# Resin acids play key roles in shaping microbial communities during degradation of spruce bark

**DOI:** 10.1101/2023.04.19.537524

**Authors:** Amanda Sörensen Ristinmaa, Albert Tafur Rangel, Alexander Idström, Sebastian Valenzuela, Eduard J. Kerkhoven, Phillip B. Pope, Merima Hasani, Johan Larsbrink

**Affiliations:** Department of Life Sciences, Chalmers University of Technology, SE-412 96 Gothenburg, Sweden; Novo Nordisk Foundation Center for Biosustainability, Chalmers University of Technology, SE-412 96 Gothenburg, Sweden; Department Chemistry and Chemical Engineering, Chalmers University of Technology, SE-412 96 Gothenburg, Sweden; Department of Medical Biochemistry and Cell Biology, University of Gothenburg, SE-405 30 Gothenburg, Sweden; Faculty of Biosciences, Norwegian University of Life Sciences, NO-1433 Ås, Norway; Faculty of Chemistry, Biotechnology and Food Science, Norwegian University of Life Sciences, NO-1433 Ås, Norway; Wallenberg Wood Science Center, Chalmers University of Technology, SE-412 96 Gothenburg, Sweden

**Keywords:** *Pseudomonas*, resin acid, abietic acid, spruce bark, metagenomics, whole genome sequencing

## Abstract

The bark is the outermost defense of trees against microbial attack, largely thanks to toxicity and prevalence of extractive compounds. Nevertheless, bark decomposes in nature, though by which species and mechanisms remains unknown. Here, we have followed the development of microbial enrichments growing on spruce bark over six months, by monitoring both chemical changes in the material and performing community and metagenomic analyses. Carbohydrate metabolism was unexpectedly limited, and instead a key activity was metabolism of extractives. Resin acid degradation was principally linked to community diversification with specific bacteria revealed to dominate the process. Metagenome-guided isolation facilitated the recovery of the dominant enrichment strain in pure culture, which represents a new species (*Pseudomonas abieticivorans* sp. nov.), that can grow on resin acids as a sole carbon source. Our results illuminate key stages in degradation of an abundant renewable resource, and how defensive extractive compounds have major roles in shaping microbiomes.

## Introduction

Bark is the outer protective barrier of trees and shields them from abiotic and biotic stress, both thanks to gradual peeling during growth and its chemical composition. The bark structure differs from wood in that it typically contains a higher amount of so-called extractive compounds (extractives), around 20-30%, in addition to the regular wood polymers cellulose, hemicelluloses, and lignin^1–3^. Extractives are secondary metabolites with diverse properties that can be extracted using hydrophilic or hydrophobic solvents. Compared to wood, the chemical composition of bark remains less well understood and it also differs widely among tree species, especially regarding the extractives, and may also differ depending on season, and sampling point^4^. Spruce is one of the dominant species and of high economic value in the northern hemisphere, and its bark which comprises around 10-15% of the harvest volume contains a high amount of lipophilic extractives, mainly resin acids (>10 mg/g dry bark), triglycerides (∼8 mg/g), steryl esters (∼5 mg/g), fatty acids (∼2.5 mg/g), and sterols (∼1.3 mg/g) (Fig. 1)^3^.

**Figure 1.**
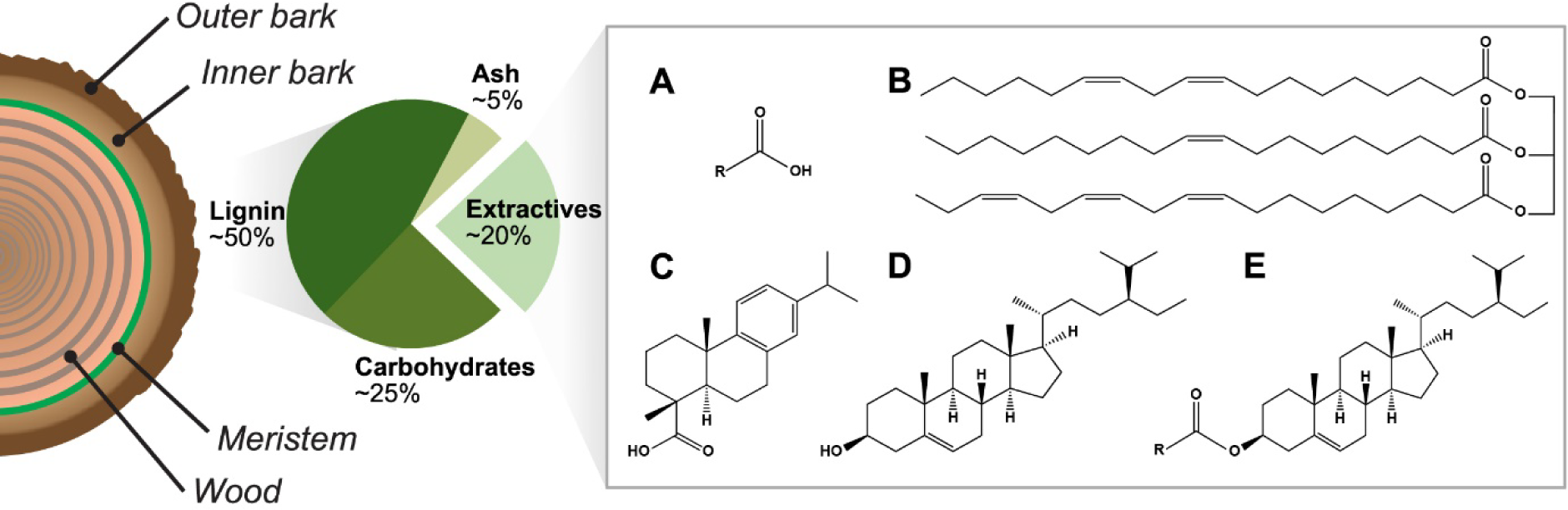
Overview of the structure of tree bark. The bark can be classified as inner or outer, which depends on its proximity to the external environment and is governed to a large extent by its age. In addition to the typical components of lignocellulose, the bark contains many additional molecule types with highly varying properties classified as extractives. Lipophilic extractives groups in spruce bark extract: A) fatty acids (R=C17, octadecanoic acid), B) triglycerides (glycerol linked to the three unsaturated fatty acids octadecenoic acid (C18:1); octadecadienoic acid (C18:2); octadecatrienoic acid (C18:3), C) resin acids (dehydroabietic acid), D) sterols (β-sitosterol), and E) steryl esters (β-sitosterol – R=C17, octadecanoic acid).

Extractives generally inhibit microbial attack through cellular toxicity or by precipitating proteins, and this is also true for spruce extractives^5, 6^. Resin acids are known to be especially toxic to water-living organisms, with LC_50_ values (concentration lethal to half of a population) of sub-mg/L^6^. Despite the antimicrobial properties of bark, it is degraded in nature over time, but compared to the extensive literature on biological degradation of wood, data on the molecular details of microbial bark degradation are sparse. Moreover, removal of extractive compounds from softwood bark has been shown to enhance its saccharification^7^. Few studies have investigated fungal degradation of bark, but include a proteomic study of the filamentous fungus *Aspergillus nidulans* on cork oak bark, though performed on a single timepoint using a 2D gel electrophoresis setup^8^.

Although overall bark degradation studies are limited, some data exist on degradation of individual spruce extractives: non-toxic triglycerides and fatty acids, as well as toxic resin acids. Triglycerides have been shown to be degraded by the fungi *Phlebiopsis gigantea, Trametes versicolor,* and *Bjerkandera spp*^9, 10^. Additionally, sterols and unsaturated fatty acids were degraded by *Phanerochaete velutina* and *Stropharia rugosoannulata*^11^, and free and esterified sterols by *Phlebia radiata* and *Poria subvermispora*^12^. Bacteria have also been shown to degrade triglycerides, sterols, and resin acids^13^. More detailed investigations of resin acid degradation by *Pseudomonas* and *Paraburkholderia* suggested a degradation pathway of abietane-type resin acids^14, 15^. However, it is presently unclear whether certain extractive groups have a larger effect on microbial communities and their development during bark degradation, and whether individual microorganisms tolerate, metabolize, and/or detoxify bark extractives. The composition and development of microbial communities during active decomposition of bark cannot be directly compared to similar studies of wood degradation, due to the large proportion of toxic extractive compounds in the bark material^16, 17^.

Here, we have investigated spruce bark degradation over six months using an enriched microbial community, using a combination of methods to obtain a broad picture of the entire process. The enrichment development was studied by marker gene and metagenomics sequencing, and the simultaneous chemical changes of the carbohydrates, lignin, and extractives were monitored. We identified resin acids as key molecules to be degraded to allow the expansion of microbial diversity, and carbohydrates were curiously only metabolized to a lesser extent. Bacteria were found to dominate the community, and from this we successfully isolated and genome-sequenced the main resin acid-degrading bacterium, representing the new species *Pseudomonas abieticivorans*. We confirmed that *P. abieticivorans* can grow on resin acids as the sole carbon source, which is attributed to a large gene cluster encoding this dedicated function. Our holistic study paves the way for detailed understanding of microbial degradation of bark, which is important information concerning both the overall carbon cycle in forests and future development of valorization strategies for complex renewable bark biomass.

## Results

To identify the microbial factors that facilitate spruce bark degradation, we sought to enrich and characterize specific populations that drive these processes. To sterilize the bark without excessively disrupting its structure, it was gamma-irradiated instead of autoclaved, as elevated temperatures and pressure would release or modify the extractives. The bark was inoculated with an actively growing inoculum prepared from an industrial bark pile and analyzed over time for chemical changes and microbial community composition (Fig. 2), up to six months (24 weeks), including an abiotic control sample (blank).

**Figure 2.**
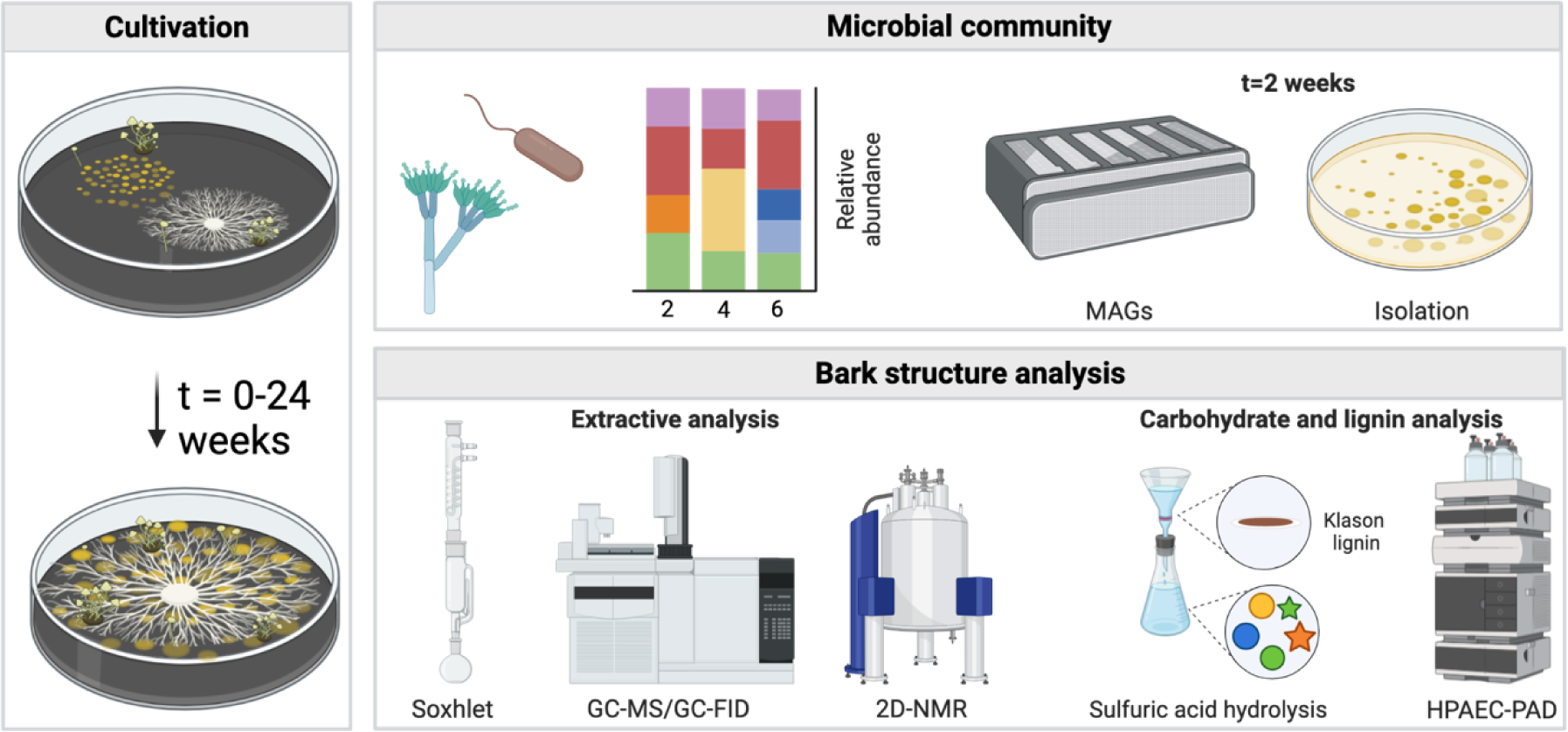
Experimental design to investigate microbial bark degradation. Community analysis (16S/ITS rRNA sequencing) was integrated with chemical analyses of the bark itself to map both community development and identify key degradative stages. In-depth metagenomics through metagenome-assembled genomes (MAGs) was performed on a chosen timepoint, from which also strain isolation was performed. Created with BioRender.com.

### Chemical analyses of microbial spruce bark degradation

To catalog the organic and inorganic landscapes of the bark throughout the six-month cultivation cycle, we first analyzed the major components of the bark: carbohydrates, lignin, extractives, and ash, for each time point. The ash content appeared to increase by approximately 53% in the biotic samples over the entire cultivation (Fig. S1), which we ascribe to loss of CO_2_ or other volatiles, and therefore it was used to normalize the data in the biotic samples. The overall mass loss represents 3.6 g per 10 g sample, but the total bark consumption should be considered higher as microbial cells also represent a fraction of the total carbon. The bark carbohydrates were expected to be a major nutrient source for the consortium, and the monosaccharide composition was determined. Surprisingly, there were only small decreases in hemicellulose-derived monosaccharides (mannose, galactose, arabinose, xylose) (Fig. S1 & Fig. S2). However, a larger effect could be seen for glucose (Fig. S1 & Fig. S2), and attributed to cellulose- or perhaps more likely starch degradation, as spruce bark can contain significant amounts of this more accessible polysaccharide^4^. Some of the measured monosaccharides likely also originate from the microorganisms, which could not be separated from the bark itself. Lignin degradation is generally a slow process performed by white-rot fungi, and potential degradation was measured by monitoring acid-insoluble and acid-soluble lignin, which showed minor differences between biotic and abiotic samples (Fig. S1 & Fig. S2). These collected results imply that the largest effect of microbial degradation could be found in the extractives.

The bark was extracted using acetone, to enable broad recovery of different extractive types, and the extractive yield of the untreated sample was similar to previous studies using acetone (Fig. S1 & Fig. S2)^4^. As hypothesized, the amount of extractives decreased significantly over time for the biotic sample with 46% decrease in extractive dry weight after two weeks and 76% decrease after 24 weeks. In the untreated bark no significant differences were observed (Fig. S2), indicating that bark extractives are a major carbon source for initial metabolism of spruce bark. Possibly, the extractives also prevent growth by specialized carbohydrate-degrading microorganisms.

### Extractive analysis reveals microbial degradation of resin acids

To investigate which extractive types were mainly degraded, they were identified and quantified using gas chromatography (GC-MS and GC-FID, Table S1), and further characterized using 2D-NMR. As mentioned, resin acids are major lipophilic extractives in spruce bark, predominantly dehydroabietic acid, abietic acid, isopimaric acid, and 7-oxodehydroabietic acid, and these are regarded as highly toxic to microorganisms^6^. Interestingly, we observed drastic decreases of the resin acids, especially during initial growth, apart from 7-oxodehydroabietic acid which was rather stable throughout the cultivation. 7-oxodehydroabietic is however a known degradation intermediate^15^, and our results suggest continuous resin acid degradation (Fig. 3). The apparent increase in dehydroabietic acid in the first time point may suggest release from the bark matrix, which we cannot explain since resin acids are not known to be linked to other bark molecules. However, the observation may point to some of the resin acids being, for instance, weakly bound to the bark matrix or abietic acid being microbially transformed into dehydroabietic acid during the cultivation^18^. Resin acids are known to be significantly affected during wood storage, though no changes were observed in the abiotic control, suggesting that observed resin acid alteration is mainly a biotic effect, in addition to known abiotic effects such as oxidation (Fig. S3)^19^. Further 2D-NMR analysis of the two-week sample confirmed dehydroabietic acid as the main extractive, with agreement both of expected aromatic and carboxylic structural signals (Fig. S4). For the later week samples, 7-oxodehydroabietic acid was the main compound (by GC) (Table S2). However, interfering signals in the sample and the relatively low concentration of 7-oxodehydroabietic acid confounded confirmation by NMR.

**Figure 3.**
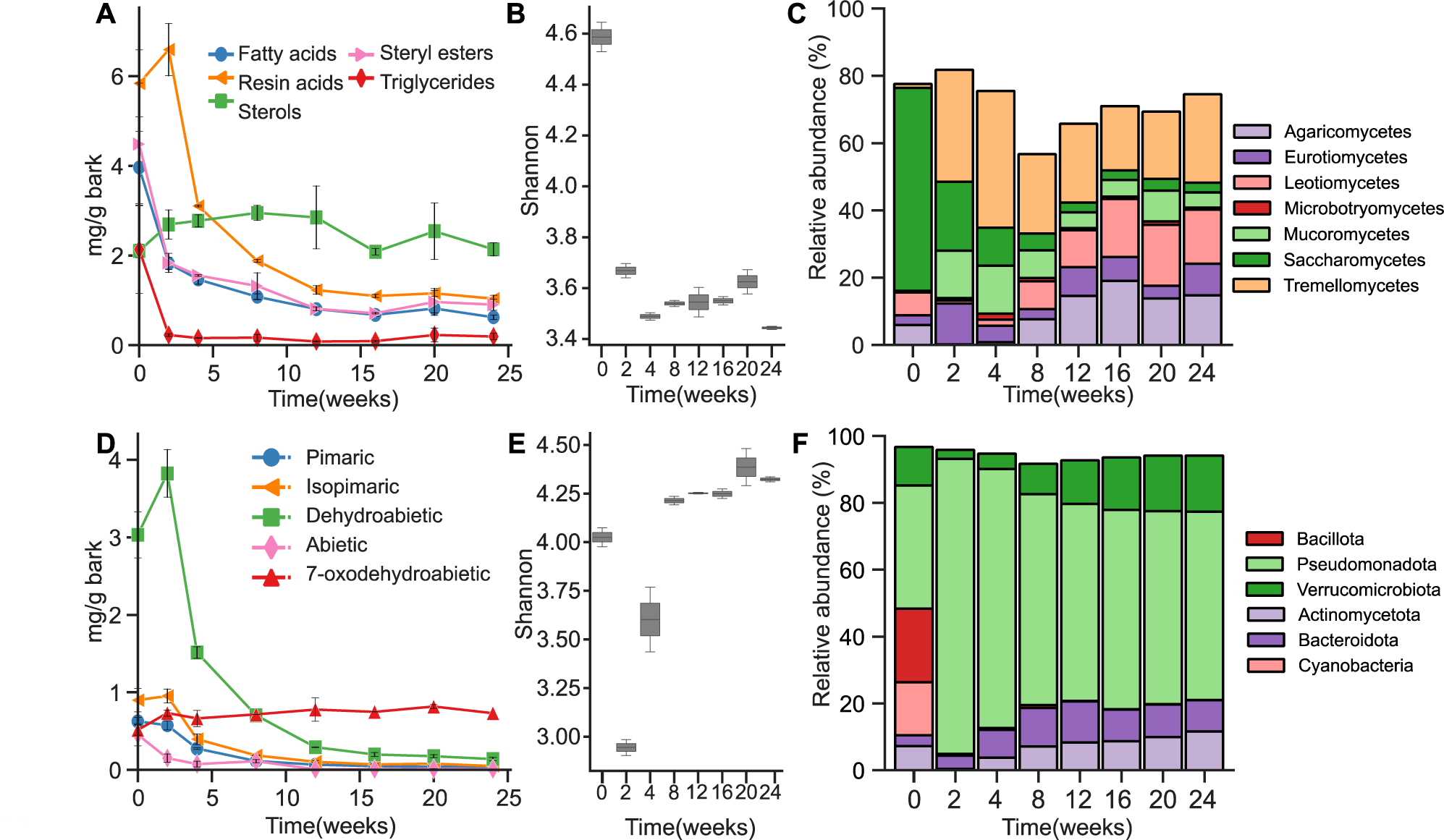
Effect of inoculating sterile bark with a culture containing a mixture of both bacteria and fungi. A) Lipophilic extractive groups in the acetone extract over time. The resin acid concentration decreased after two weeks, and the sterol concentration increased over time. D) Individual resin acid compounds, which all decreased over time except 7-oxodehydroabietic acid. Alpha diversity of the B) fungal community, and E) bacterial community over time, expressed as the Shannon diversity index. C) Relative abundance of the different classes in the fungal community over time derived from two biological replicate experiments. F) Relative abundance of the phyla in the bacterial community over time. Mean values and standard deviations are based upon duplicate biological experiments.

The majority of fatty acids in spruce bark are oleic and pinoleic acids, while nearly all sterol and steryl esters are β-sitosterol and β-sitosterol linked to fatty acids^20^. Of these non-toxic compounds, the triglycerides were rapidly metabolized and almost entirely consumed after two weeks, while for fatty acids and steryl esters there was a sharp initial drop followed by a gradual decrease (Fig. 3). Such degradation has previously been shown for both fungi and bacteria^9, 13^. Again, no abiotic effects were observed, such as spontaneous hydrolysis of steryl esters (Fig. S3). The sterols in the actively growing culture increased initially (weeks 0-8), and then instead decreased (weeks 8-24), which could both be derived from the microbial growth (membrane sterols) or release from steryl esters. The long-chain fatty acids and alcohols all decreased over time, though in the blank sample they increased, possibly due to spontaneous oxidation (Fig. S3). Additionally, a ferulic ester compound could be identified in the later timepoints using different 2D-NMR methods, where the aromatic region as well as the ester bond were confirmed rather than a free ferulic acid (Fig. S5). Possibly, this ester is derived from partial degradation of suberin.

### Enrichment of resin acid degrading bacteria in spruce bark microbial communities

To compare the chemical changes in the bark to the development of the microbial consortium enriched on it, we analyzed its composition by sequencing of marker gene regions of both bacteria (V4/16S rRNA) and fungi (ITS2), followed by taxonomical assignment, and a principal component analysis (PCA) showed good agreement between replicates (Fig. S6). After quality control and bioinformatic processing between 19,047 and 29,052 fungal reads (Table S3), and 38,037 and 70,628 bacterial reads (Table S4) were generated per sample. The alpha diversity of the fungal community drastically decreased already at the first timepoint and remained low relative to the inoculum, indicating a strong selection pressure by the bark (Fig. 3 & Fig. S7). During early colonization Saccharomycetes drastically decreased in abundance compared to the original inoculum, whereas Mucoromycetes and especially Tremellomycetes increased. Interestingly, both Agaricomycetes and Leotiomycetes were below detection levels in the first timepoint but then reappeared after four weeks and gradually increased in proportion, suggesting they are completely inhibited by the extractives, or possibly by the species that metabolize these. In the two-week sample, the fungal community was dominated by *Apiotrichum* (32.8%), *Mucor* (14.1%), *Trichoderma* (6.55%), and *Penicillum* (11.55%), which are common cosmopolitan fungal genera.

Similar to the fungal community the diversity of the bacterial community sharply decreased in the two-week sample. However, it contrastingly increased thereafter and reached an even greater diversity than the inoculum in the eight-week sample (Fig. 3 & Fig. S7). This could reflect the generally quicker growth rate of bacteria, or greater abilities to metabolize the inhibitory bark substances. During early colonization the bacterial culture was dominated by populations affiliated to Bacillota (Firmicutes) and Cyanobacteria, which virtually disappear after two weeks. Phyla that quickly colonized the bark were Pseudomonadota (Proteobacteria), and Bacteroidota (Bacteroidetes), However, while Pseudomonadota decreased in relative abundance after initial colonization, they remained dominant. Interestingly, after 8-12 weeks the relative abundances of Bacteroidota and Verrucomicrobiota had increased to a sizable fraction, which could indicate a shift toward carbohydrate utilization as these are phyla known to comprise carbohydrate specialists^21, 22^. Overall, the bacterial microbial community was dominated by the genera *Pseudomonas*, *Burkholderia-Caballeonia-Paraburkholderia*, and *Paravibacterium* which are known to be fast-growing. To gain further insight into the microbiome at two weeks this sample was further sequenced to reconstruct metagenome-assembled genomes (MAGs).

### Metagenomics divulges potential key resin acid-degrading bacteria

The combined observation of resin acid degradation and low diversity during initial growth suggests enrichment of highly specialized microorganisms that can tolerate, detoxify, or fully metabolize resin acids. To gain deeper biological insight on the process, shotgun metagenome sequencing was performed on the 2-week sample and metagenome-assembled genomes (MAGs) reconstructed (Table S5). The taxonomic structure of the metagenomics confirmed bacteria as being dominant (76% of reads/contigs) relative to eukaryotes (8%), and a portion of unclassified reads (18%) (Fig. S8). 15 MAGs were binned resulting in 13 MAGs of high quality (>90% completion, <5% contamination) and two of medium quality with more than 5% contamination or no 16S rRNA (Table S6). As in the 16S rRNA analyses, the metagenome (MG) showed that Pseudomonadota were most abundant and comprised of Alphaproteobacteria (37%), Betaproteobacteria (13%), and Gammaproteobacteria (11%) (Fig. S8), representing either unknown species or genera including known resin acid degraders such as *Pseudomonas* and *Burkholderia*^15, 18^.

Resin acid-degradation is sparsely described in literature but within the genomes of *Pseudomonas abietaniphila* BKME-9, *Pseudomonas diterpeniphila* A19*-*6a, and *Paraburkholderia xenovorans* LB400 (formerly: *Burkholderia xenovorans*) a so-called *dit* gene cluster has been attributed to this function from gene knockout studies (Table S7)^15, 23, 24^. The presence of genes homologous to *dit* genes from *P. abietaniphila* and *P. xenovorans* in the MAGs was investigated (Fig. 4 & Fig. S9), and a complete *dit* cluster was identified in MAG15 which was also the most abundant microorganism (19.2%) in the metagenome (Fig. 4). Varying *dit* cluster completeness was found in the other MAGs from BLAST comparisons (≥30% seq. id.), indicating that these organisms have either incomplete resin acid degradation pathways, unknown pathways, or are unable to fully metabolize resin acids. Many of the MAGs classified as Pseudomonadota (MAG 1, 4, 7, 9, 14, 15) have a *dit*A1 homolog which is a gene shown to be essential for resin acid-degradation through gene knockout analyses in both *P. abietaniphila* and *P. xenovorans*^15, 23^. In the metagenome, MAG7 was the second most abundant (12.6%) and classified as *Paraburkholderia*. It encodes a putative partial *dit* cluster comprising *dit*A1, *dit*B, *dit*D, *dit*I and *dit*Q (Fig. S10). Despite MAG14 being third most abundant and identified as *Pseudomonas*, the only putative co-located *dit* genes are a homolog of *dit*A1 and *dit*B interspersed with two genes of unknown function (Fig. S10). MAG6 was identified as *Paraburkholderia tropica* with an ANI score of 99.06%, which despite only being 4.3% relative abundant has a considerable amount of putative *dit* genes, including two *dit*A1 genes found in two different clusters, together with *dit*F and *dit*J homologs. Curiously, MAG4 encodes a seven-gene cluster comprising *dit*A1, *dit*A2, *dit*C (24.5% seq. id.), *dit*B, *dit*I, *dit*L and *dit*Q, while only representing 2% of the metagenome reads.

**Figure 4.**
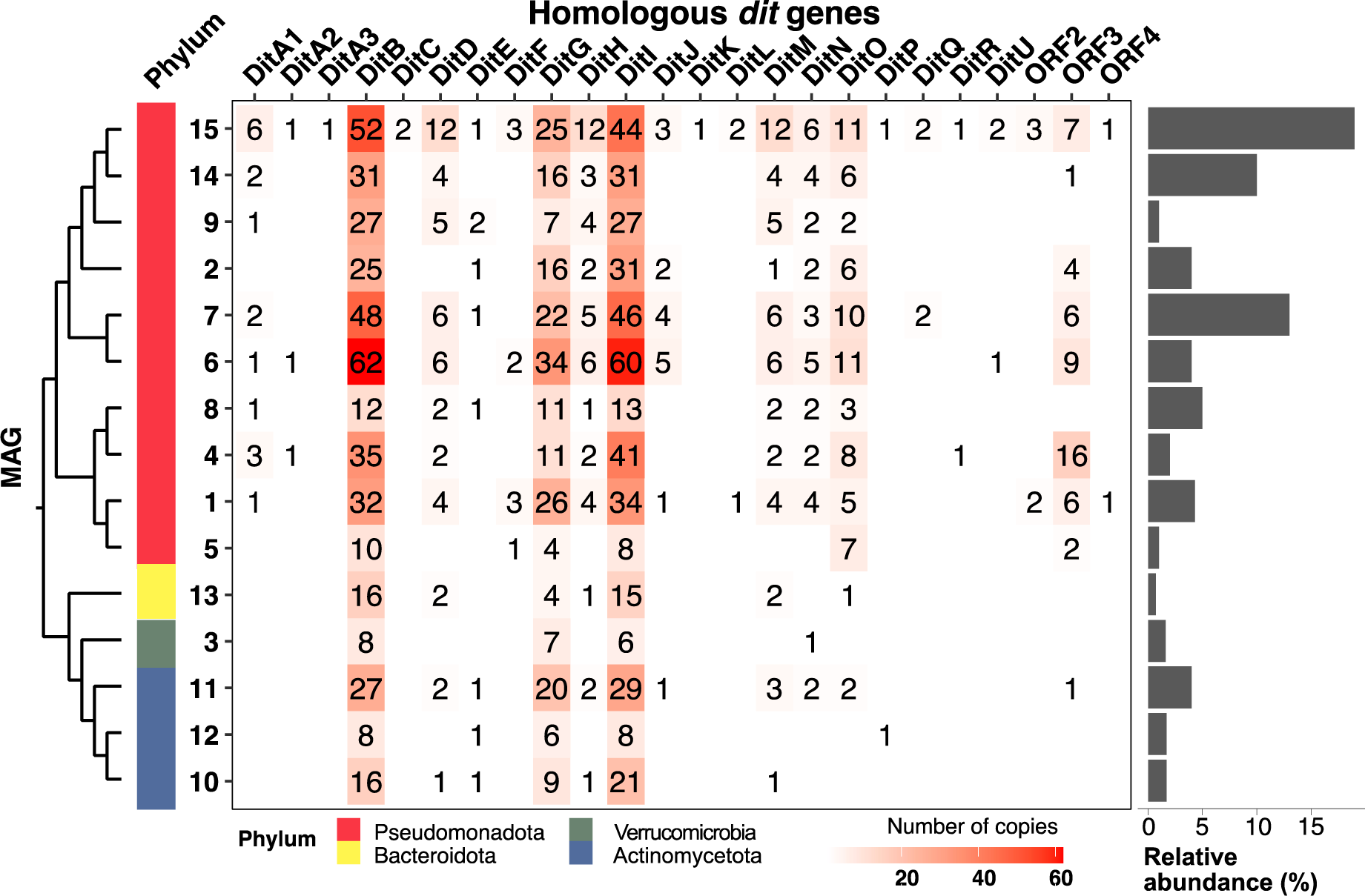
Metagenome Assembled Genomes (MAGs) from the two-week spruce bark degradation sample. Proteins were identified in the MAG sequences by comparing with the *dit* clusters in *P. abietaniphila* BKME-9 and *P. xenovorans* LB400 using protein-protein BLAST with a E-value of 1e^−10^ and sequence identity ≥30%, and duplicate sequences with the same Dit protein hit and query were removed. Partial gene clusters (co-located genes) are found in Fig. S10. Taxonomic classification is based on GTDB (Genome Taxonomy Database), and phyla indicated by color^28^. MAGs are clustered according to proteome-based GBDP distance (Genome Blast Distance Phylogeny) using the TYGS database^29^. This type of phylogenetic clustering was used as no 16S rRNA sequence could be extracted from MAG12. The relative abundance of MAGs (determined via CoverM v. 0.6.1.) within the metagenome is shown on the right.

While extractive degradation was the main activity of the community, we sought to identify potential carbohydrate-active enzymes (CAZymes) present in the MAGs using dbCAN2^25^. In total, 1783 putative CAZymes were identified, including 777 glycoside hydrolases (GHs), 752 glycosyltransferases (GTs), 135 carbohydrate esterases (CEs), 81 auxiliary activities (AAs), and 38 polysaccharide lyases (PLs) (Fig. S11). Interestingly, the lowest abundant MAGs encoded the highest number of degradative CAZymes (non-GT), supporting our chemical analysis results that imply that carbohydrates are not the main nutrient during initial bark degradation (Fig. S11-S15). MAG13 (0.7% relative abundance) stood out as a likely polysaccharide specialist from the Bacteroidota, and it encodes 115 putative GHs, many of which are found in putative polysaccharide utilization loci (PULs) – large gene clusters comprising sugar capture-, transport-, degradation-, and regulatory proteins^22^. MAG13 encodes 37 of the PUL signature SusC/SusD-like pairs (sugar transport/capture), in ten different PULs (Fig. S16). Two of these PULs appear to target pectin (rhamnogalacturonan I side chains and rhamnogalacturonan II, respectively), two hemicelluloses (xyloglucan and galactoglucomannan, respectively), and one starch (Fig. S16), which are all polysaccharides found in spruce bark^4, 26, 27^. The other PUL targets are not easily identified but could be extracellular polymeric substances (EPS) from competing microorganisms.

### Isolation, genome sequencing, and phenotyping of the dominant resin acid-degrading bacterium

With resin acid degradation being a major activity in early spruce bark degradation and MAG15 being numerically dominant, we pursued to study this specific population in more detail. Based on the previous chemical and genomic information, we designed culture conditions with abietic acid as a carbon source which enabled us to successfully isolate a bacterial strain (designated PIA16) from the two-week sample which was further characterized via genome-sequencing and phenotypic analysis. Long sequencing reads were assembled into a reference-grade, closed 6,715,763 bp genome, with a 62% G+C content and 5,987 predicted protein-coding genes (Table S8 and Table S9). A significant Average Nucleotide Identity (ANI) match (>99%) between the isolate and MAG15 supported it being the same species (Fig. S17), and affiliated to the *Pseudomonas* genus, while taxonomic classification analyses using both the Genome Taxonomy^29^ and SILVA^30^ databases supported our strain being a new species, which we named *Pseudomonas abieticivorans* sp. nov.. Sequence comparisons of 16S rRNA sequences of *P. abieticivorans* and 181 other *Pseudomonas* species (total 269 sequences) showed *P. abieticivorans* being located on a separate phylogenetic branch (Fig. S18), supporting its classification as a new species. ANI analysis showed the most closely related species being *P. baetica* a390 (81.4%), *P. putida* NBRC 14164 (81.6%), and *P. koreensis* Ps 9-14 (81.6%) (Table S10), much lower than the recommended 95% cut-off to distinguish species. Differential phenotypic tests compared to closely related strains are shown in Table S10, and further differentiate *P. abieticivorans* biochemically from these. The ANI similarity between *P. abieticivorans* and species previously shown to degrade resin acids was also low, with values of 81.9%, 80.6%, and 80.15% to *P. vancouverensis, P. abietaniphila,* and *P. multiresinivorans*, respectively. The final growth density of *P. abieticivorans* on resin acids was observed to be fairly low (maximum OD_600_ ∼0.2), and it is interesting to speculate what limits growth on these substrates, but as the solubility of resin acids is very low (1.7-5 mg/L), solubilization is likely a major factor limiting utilization^6^.

Genome analysis of *P. abieticivorans* revealed few degradative CAZymes, and only suggests the ability to grow on starch, with enzymes from GH13 (7 homologs), GH31 (1), GH77 (1), in addition to several putative enzymes involved in bacterial cell wall lysis/remodeling (e.g., GH23 (5), GH73 (2), GH103 (2)) (Fig. S19). The API ZYM enzyme activity screen showed a negative result for β-galactosidase and β-glucosidase, but positive for esterase lipase (C8) and lipase (C14) (Table S10), which further supports the hypothesis that this species does not primarily consume polysaccharides but rather extractive compounds in spruce bark. Similar to other resin acid degrading isolates, *P. abieticivorans* can grow on both dehydroabietic and abietic acid as a sole carbon source (Fig. 5), but interestingly it can also grow on the more toxic isopimaric acid^6^, which is an unusual trait (Fig. S20)^18^. To gain insight into correlations in protein coding sequences between known resin acid degrading strains we performed a genome collinearity analysis (Table S11), identifying 15,765 (40.67%) sequences involved in 639 synteny blocks (Fig. S21), defined as a block of five orthologous genes found in the same order between the compared genomes^31^. As expected, fewer collinearity blocks were observed between *P. abieticivorans* and *Paraburkholderia xenovorans* compared to *Pseduomonas* strains. Interestingly, when analyzing the inter-species synteny blocks, it was observed that *P. abieticivorans* shares a high collinearity level in two regions (bases 5,144,228 to 5,171,342, and 6,497,666 to 6,541,998) which is composed of around 80 protein-coding genes (Fig. S21). When analyzing the number of times that a gene was included inside a synteny block and shared among species, it was observed that region 0.7 – 4.7 Mb in the genome of *P. abieticivorans* are frequently shared between *Pseudomonas sp.* genus studied here. One such shared synteny block contained clusters of genes previously identified to sustain growth on resin acids (Fig. S21).

**Figure 5.**
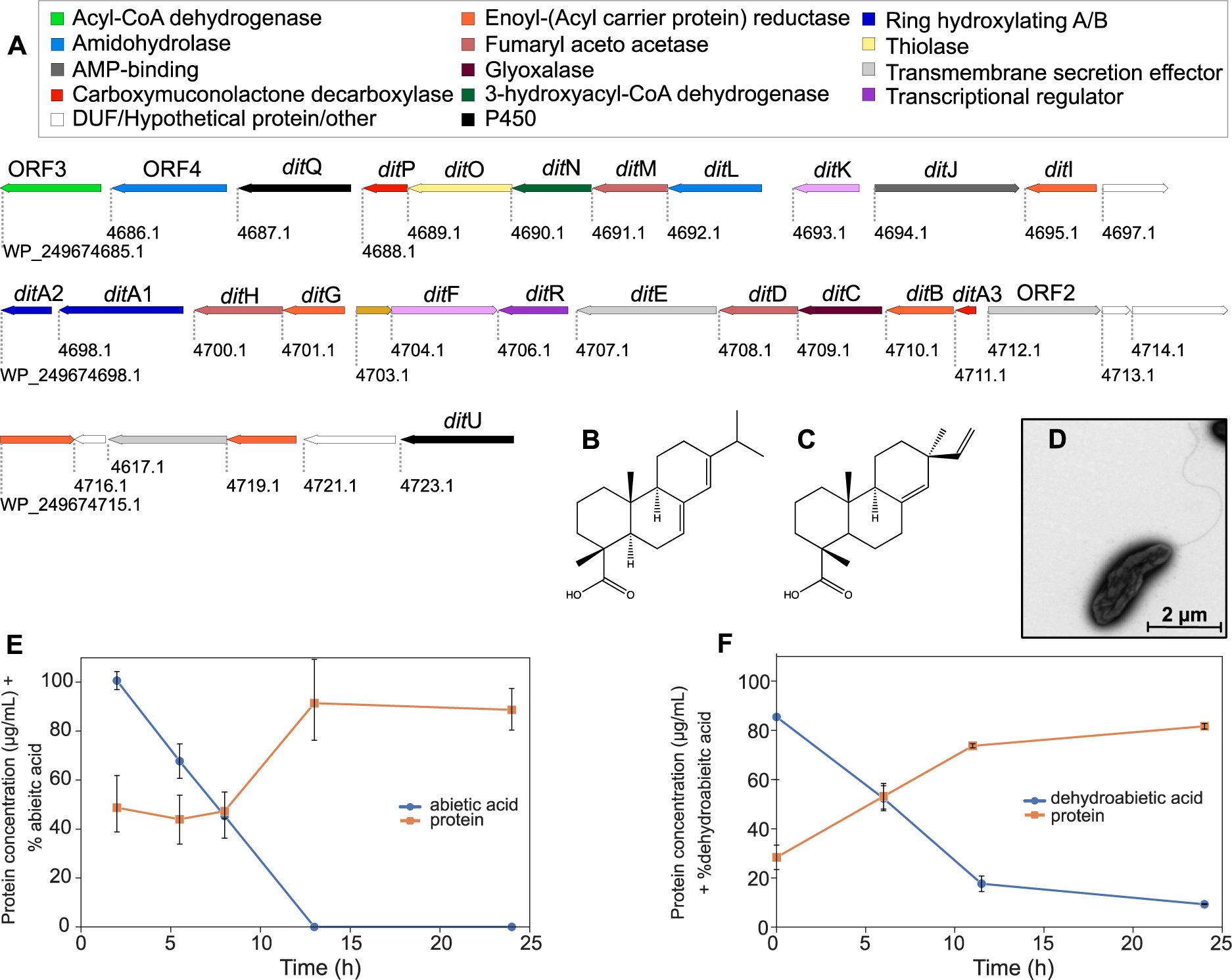
The genetic organization of the *Pseudomonas abieticivorans dit* cluster and growth on resin acids. A) The identified *dit* cluster which spans 34.6 kbp. Genbank accession numbers are shown below each gene, gene names above, and their functional annotations according to pfam are color coded. The resin acids are divided into two subtypes: abietane and pimarane, the former represented by B) abietic acid, and the latter by C) isopimaric acid. D) Representative transmission electron microscopy image of a *P. abieticivorans* cell, showing the presence of a single flagellum. Degradation of resin acids by *P. abieticivorans* on E) abietic acid and F) dehydroabietic acid, and concomitant cellular protein concentrations as a proxy for growth as optical density monitoring is complicated by the insoluble nature of the resin acids. The means and standard deviations are based on duplicate biological experiments.

As expected from the metagenomics, *P. abieticivorans* encodes a complete *dit* cluster (24/24 expected genes), which was compared to those found in known resin acid-degrading strains using BLAST and collinearity analyses (Fig. 5). Strikingly, *P. abieticivorans* had an identical *dit* cluster genetic organization as that of *P. abietaniphila*, though only partial overlap was observed with *P. xenovorans* (Fig. S21). A lower number of co-located *dit* genes (10/24) were also identified in *P. resinovorans*, but in *P. multiresinovorans*, homologs to *dit* genes were not found in a cluster but rather spread out in the genome.

## Discussion

Bark is vital for the protection of trees and disruption of this outer barrier by e.g., wood-boring insects can lead to enhanced microbial access, decimated coniferous forests worldwide and disruption of global ecosystems^32, 33^. Despite its fundamental protective role, how the bark is degraded by microorganisms is not known. Previous studies have shown that microbial communities differ between bark and wood, as well as the specific sampling point on the tree, and also that the presence of bark slows down the degradation of the underlying wood^34^. The importance of the bacterial community on bark was recently highlighted where it was shown that bark-dwelling methanotrophic bacteria decrease methane emissions from trees, thus mitigating global warming^35^. Fungal bark communities have been less investigated but shown to change over time as wood degrades through measurements of wood decay stage, density, moisture, and C:N ratio^36^. Despite the importance of extractives in the antimicrobial properties of bark, they have however previously been overlooked.

In this study, we monitored the development and dynamics of a bark-sourced microbial community on fresh spruce bark over six months, as well as the chemical changes to the material to identify important degradation stages, compounds, and microorganisms for this process. The minor degradation of polysaccharides was surprising, as especially non-cellulosic and non-crystalline carbohydrates are regarded as readily degradable by plant biomass-metabolizing enzymes, while slow lignin degradation was observed as expected. Instead, it appears that for the enrichments used in this study, the bark extractives are the main nutrients in the initial degradation stages. The correlation between the drastic decrease in resin acid concentration in the bark and the sharp decrease in alpha-diversity in both the fungal and bacterial communities was pronounced and suggestive that resin acids may act as ‘gate-keeper’ molecules and that their detoxification is a key step in bark degradation. This agrees well with resin acids being regarded as major defensive compounds with broad antimicrobial properties^37^.

Resin acids are thought to mainly be encapsulated within resin canals in wood^38^, and the apparent increase in the concentration of especially dehydroabietic acid at the initial degradation stage is interesting. It could indicate that the resin acids are in fact inaccessible in fresh bark and possibly weakly bound to other molecules, polymerized, or otherwise tightly encapsulated within the bark and before being released. Currently, there is no information on resin acids being for instance ester-bonded to any structural component within the bark, but it could be speculated that their release is instead linked to the degradation of triglycerides, as these can help solubilize the hydrophobic resin acids in the resin itself. Recently, extractive compounds have been shown to either inhibit or enhance the activity of carbohydrate-active enzymes, such as lytic polysaccharide monooxygenases (LPMOs)^39^, so it is possible that the extractives may exert general enzyme-inhibitory rather than toxic effects that block wider carbohydrate utilization.

Our deeper metagenomic analysis revealed the abundance of specialized MAGs and their *dit* cluster-related genes, assumed to encode enzymes and proteins involved in resin acid (diterpene) degradation^15, 18, 23^, which further indicates that resin acid metabolism is crucial to thrive on fresh spruce bark. Interestingly, we could observe an inverse relationship with encoded degradative CAZymes in our reconstructed MAGs, where populations with higher MAG abundance encoded smaller CAZyme profiles and vice versa (Fig. 4 & Fig. S11). Over a longer time frame, the amplicon data suggest that the bacterial populations begin shifting from an extractive-focus to a carbohydrate conversion, given the increase of Bacteroidota and Verrucomicrobiota^21, 22^, though a clear shift in actual carbohydrate-degrading capacity in the enrichment did not become apparent during our study. Our metagenomic analysis also facilitated the discovery and subsequent characterization of *P. abieticivorans*, which dominated our enrichment cultures and encodes a complete *dit* cluster. The molecular basis for resin acid degradation has not been demonstrated apart from one study of a cytochrome P450 monooxygenase from *Streptomyces griseolus*^40^, though whether resin acids are a natural substrate for this enzyme is unclear. Further investigation of *P. abieticivorans* and related species can help shed light on this important process, which likely involves solubilization of the resin acids at some stage to facilitate import into the cell.

Overall, our study indicates that microbial degradation of the bark is a slow process, and also that abiotic changes to the material are minor even over our six-month experiment. Spruce is indispensable to the forest industry and is used for pulp and paper production in the northern hemisphere. Looking only at the Nordic countries, bark is an industrial side stream produced in close to 400 million m^3^ per year^41^, which today is poorly utilized. Understanding how it can be degraded in nature by different microorganisms is key for future valorization, for instance using select enzymes, and can also be of high relevance to limit the harmful effects of wood-boring insects. Bark from other species contain other types of extractives, with for example birch bark being enriched in betulin and pine in condensed tannins^41^, and our study sets a basis for investigating whether degradation also of other bark types progresses via initial degradation of key ‘gatekeeper’ molecules that strongly inhibit the growth of but a few extractive-detoxifying microbial species.

### Description of Pseudomonas abieticivorans sp. nov

*Pseudomonas abieticivorans* (a.bi.e.ti.ci.vo’rans. N.L. neut. n. acidum abieticum, abietic acid; L. pres. part. vorans, eating; N.L. part. adj. abieticivorans, eating abietic acid). Cells are Gram-negative, rod-shaped (0.77 μm wide, 2.6 μm long), with a singular flagellum, and catalase and oxidase negative. Growth occurs at 4-30 °C. Optimal growth temperature is between 25-28 °C. The species grows at pH 5-9 and in the presence of 6 w/v NaCl. The species is negative for reduction of nitrates to nitrate and reduction of nitrates to nitrogen, for indole production, fermentation of glucose, urease, and proteases for gelative hydrolysis. The species is positive for assimilation of D-glucose, L-arabinose, D-mannitol, potassium gluconate, capric acid, malate, and trisodium citrate, and negative for assimilation of D-mannose, *N*-acetyl-D-glucosamine, D-maltose, adipic acid, and phenylacetic acid. The enzymatic activities which were positive were alkaline phosphatase, esterase (C4), esterase lipase (C8), leucine arylamidase, valine arylamidase, naphthol-AS-BI-phosphohydrolase, acid-phosphatase, arginine dihydrolase, and catalase. The ones which displayed a negative test were lipase (C14), cystine arylamidase, trypsin, α-chymotrypsin, β-galactosidase, β-glucuronidase, α-glucosidase, β-glucosidase, *N*-acetyl-glucosaminidase, α-mannosidase, α-fucosidase. Growth is detected in R2A, PIA, and LB. The major fatty acids are summed feature 3 (C_16:1_ ω7c and/or C_16:1_ ω6c, 26.12%), summed feature 8 (C_18:1_ ω6c and/or C_18:1_ ω7c, 13.96%), C_16:0_ (31.57%), C_17:0_ cyclo (12.79%), C_12:0_ (4.11%), C_12:0_ 2-OH (3.05%), C_12:0_ 3-OH (3.49%), C_10:0_ 3-OH (3.00%), C_14:0_ (0.47%). The type strain, PIA16 (=CCUG 76343T, =DSM 114633) was isolated from spruce bark on abietic acid. The G+C content of the type strain is 61.1 mol% and the genome size is 6,715,763 bp.

## Methods

### Bark preparation

Spruce bark was obtained from the Iggesund pulp and paper mill (Iggesund, Holmen AB, Sweden). The bark was left to dry at room temperature for seven days. Thereafter, the bark was milled in a Wiley-type mill, mesh size <0.1 mm, and was afterwards sterilized with 25 kGy by gamma-irradiation (Mediscan GmbH & Co KG).

### Microbial cultivation

To mimic growth conditions on spruce bark in nature, solid-state cultivation was used. First, a bark inoculum was generated by growing unsterilized bark in 2.3 mL M9 minimal medium (disodium hydrogen phosphate 6.778 g/L, potassium phosphate 5 g/L, ammonium chloride 1 g/L, sodium chloride 0.5 g/L, magnesium sulfate 0.24 g/L, calcium dichloride 0.012 g/L) per g bark for six months, to obtain an actively growing culture (evaluated from visual growth). 1 g (wet weight) of the approximately 30 g growing culture was added to 10 g sterile bark with 2.3 mL M9 medium per g bark. The solid-state cultures were incubated aerobically at room temperature (∼21 °C) and samples collected after 0, 2, 4, 8, 12, 16, 20, and 24 weeks of growth in biological triplicate experiments. Approximately 4 g bark samples (dry weight) were weighed and used for chemical analysis of the carbohydrates, lignin, and extractives present in the bark, while the remaining culture was frozen at −20 °C and used for 16S rRNA and ITS profiling (Fig. 2). A non-inoculated blank sample was used as control for each time point.

### Soxhlet extraction

For evaluation of the effect of microbial growth on the spruce bark extractives, Soxhlet extraction was used. The bark samples were extracted with 200 mL acetone as solvent, with 10 min per extraction for 24 cycles. After extraction, the acetone was evaporated using rotary evaporation until 50 mL solvent remained, then left in a fume hood for evaporation of remaining solvent. The percentage of acetone-soluble extractives was determined gravimetrically based on the weight of the starting material.

### Ash content

For evaluation of the effect of microbial growth on the ash content present in the bark, the ash concentration was measured according to National Renewable Energy Laboratory standard method^42^. In brief, the weight of the inorganic residue left after 1 g of bark sample (after drying at 105 °C overnight) had been subjected to dry oxidation at 575 °C was measured.

### Chemical characterization of carbohydrates and lignin

To elucidate if the microorganisms were degrading carbohydrates or lignin in the spruce bark, the acetone-extracted bark was dried overnight, followed by hydrolysis using 72% sulfuric acid according to established protocols^43^. The insoluble residue was determined gravimetrically. The filtrate from the hydrolysis was used to determine monosaccharide and acid-soluble lignin content. The acid-soluble lignin was determined by absorbance measurements at 205 nm on a Specord 205 (Analytic Jena), using an extinction coefficient of 110 dm^3^g^−1^cm^−1^. The monosaccharide composition of the bark was monitored using high-performance anion-exchange chromatography with pulsed amperometric detection (HPAEC-PAD) on an IC5000 system (Thermo Scientific). A 2 x 250 mm Dionex Carbopac PA1 column (Thermo Scientific) with a 2 x 50 mm guard column (Thermo Scientific) was used at a column temperature of 30 °C. The eluents were A – water, B – 200 mM NaOH, and C – 200 mM NaOH and 170 mM sodium acetate^44^. The samples were eluted using eluent A (0.26 mL/min) and detected using a post-column addition of eluent B (0.13 mL/min). Thereafter, the column was washed with eluent C and equilibrated with eluent A. The monosaccharide concentrations were determined using an internal fucose standard and pure external standards of arabinose, mannose, galactose, glucose, and xylose. Peak analysis was performed using Chromeleon software 7.2.10 (Thermo Scientific).

### Extractive group analysis using GC-FID and GC-MS

To determine the effect of microbial growth on the lipophilic extractive groups present in the spruce bark acetone extracts, the bark extractives were analyzed by gas chromatography (GC) coupled to a flame ionization detector (FID) and GC-mass spectrometry (MS) by MoRe Research Örnsköldsvik AB. The bark extracts were derivatized and analyzed as trimethylsilyl derivatives. The internal standards (heneicosanoic acid, cholesterol heptadecanoate, and 1,3-dipalmitoyl-2-oleyl glycerol), at a concentration of 200 µg/mL, were added to 50 mL samples containing 30 mg/mL of bark extractives dissolved in acetone. Thereafter, the vials were evaporated under nitrogen and derivatized using 100 µL *N*,*O*-Bis(trimethylsilyl)trifluoroacetamide (BSTFA), 50 µL trimethylchlorosilane (TMCS), 20 µL pyridine, and heated for 20 min at 70 °C.

The concentrations of extractive groups (fatty acids, resin acids, sterols, steryl esters, and triglycerides) were evaluated on a Thermo Trace 1300 chromatography system coupled to an FID using an on-column Programmable Temperature Vaporization (PTV) injection method. The temperature program of the oven started at 100 °C, which was held for 1.5 min, followed by increasing the temperature to 340 °C using a gradient of 10 °C/min, and holding this temperature for 3 min. The PTV temperature increased from 80 °C to 340 °C at 10 °C/min, which was held for 10 min. The fatty acids, resin acid, and sterol ester concentrations were evaluated on a HP-5ms (30 m, 0.25 mm internal diameter, 0.25 µm film thickness) column relative to the peak area of the internal standard heptadecanoic acid. High boiling compounds, such as steryl ester and triglyceride concentrations were evaluated using a HP-1 SIM/Dist (5 m, 0.53 mm internal diameter, 0.15 µm film thickness) column based on the relative peak area towards the internal standards cholesteryl heptadecanoate and 1,3-dipalmitoyl-2-oleylglycerol, respectively. Peak evaluation was performed using Chromeleon software (Thermo Fisher Scientific).

GC-MS was used for the identification of individual compounds in the fatty acids, resin acids, and sterols present in the bark extracts. Quantification of the individual compounds was based on the FID total ion peak, relative to the peak area of the internal standard heptadecanoic acid. GC-MS analysis was performed on an Agilent 8890 chromatography system with a CTC Analytics CombiPAL (Autosampler) coupled to an Agilent 5975C MSD mass spectrometer with split-less injection using the same column and as the long column FID experiment, a HP-5ms (30 m, 0.25 mm internal diameter, 0.25 µm film thickness) column and helium as a carrier gas at a rate of 1.2 mL/min. The temperature program of the oven started at 100 °C, was held for 1 min, thereafter, the temperature was increased to 340 °C at 10 °C/min, which at the end was held for 3 min. The injector was kept at 230 °C. Compounds were identified in the MS by comparing their mass spectra with those of the Wiley and NIST libraries on MassHunter (Agilent).

### Extractive analysis using NMR

All NMR measurements were conducted on a Varian Inova 500 MHz operating at 11.7 T with a 5 mm HFX-probe. Measurements were performed at 298K with acetone-d6 as solvent. The ^1^H-measurements were conducted using an 8 µs ^1^H-detection pulse, 2 s acquisition time, 5 s recycle delay, and 32 scans. ^13^C-meaurements were conducted using a 14 µs ^13^C-detection pulse, 1 s acquisition time, 5 s recycle delay, and 8096 scans. Correlation Spectroscopy (COSY) was recorded using 2048 scans with 256 increments. Heteronuclear single quantum coherence (HSQC) was recorded using 64 scans with 384 increments and a transfer delay of 3.425 ms corresponding to a 145 Hz J-coupling. Heteronuclear multiple bond coherence (HMBC) was recorded using 64 scans with 512 increments and a transfer delay of 62.5 ms, corresponding to an 8 Hz J-coupling. For all samples, the solvent was used as chemical shift reference with the acetone-d6 methyl signal referenced to 2.05 ppm and 29.84 ppm for ^1^H and ^13^C, respectively.

### DNA extraction for fungal and bacterial gene regions

DNA extraction from spruce bark and community analysis was performed by DNAsense ApS (Denmark) using the standard protocol for FastDNA Spin kit for Soil (MP Biomedicals) with the following exceptions: 500 mL of sample, 480 mL sodium phosphate buffer and 120 mL MT Buffer were added to a Lysing Matrix E tube, and bead beating was performed at 6 m/s for 4×40s^45^. Gel electrophoresis using Tapestation 2200 and Genomic DNA screentapes (Agilent) was used to validate product size and purity of a subset of DNA extracts. DNA concentration and purity was measured using Qubit dsDNA HS/BR Assay kit (Thermo Fisher Scientific).

### Library preparation and sequencing of fungal and bacterial marker gene regions

Fungal ITS region 2 sequencing libraries and for Archaea and Bacteria, 16S rRNA gene region V4 sequencing libraries, were prepared by a custom protocol based on an Illumina protocol^46^, and performed by DNAsense ApS. Up to 10 ng of extracted DNA was used as template for PCR amplification of the Archaea and Bacteria, 16S rRNA gene region V4 amplicons. Each 25 mL PCR reaction contained 12.5 mL PCRBIO Ultra mix (PCR Biosystems) and 400 nM of each forward and reverse tailed primer mix. PCR was conducted with the following program: initial denaturation at 95 °C for 2 min, 30 cycles of amplification (95 °C for 15 s, 55 °C for 15 s, 72 °C for 50 s), and final elongation at 72 °C for 5 min. Duplicate PCR reactions were performed for each sample and the duplicates were pooled after PCR. The forward and reverse tailed primers were designed according to ^46^ and contained primers targeting the Fungal ITS region 2: ITS7 (5’-GTGARTCATCRARTYTTTG-3’) and ITS4 (5’-TCCTSCGCTTATTGATATGC-3’)^47^ and for Archaea and Bacteria, 16S rRNA gene region V4: 515FB (5’-GTGYCAGCMGCCGCGGTAA-3’) and 806RB (5’-GGACTACNVGGGTWTCTAAT-3’)^48^. The primer tails enabled attachment of Illumina Nextera adaptors necessary for sequencing in a subsequent PCR. The resulting amplicon libraries were purified using the standard protocol for Agencourt Ampure XP Beads (Beckman Coulter) with a bead to sample ratio of 4:5. DNA was eluted in 25 mL of nuclease-free water (Qiagen). DNA concentration was measured using Qubit dsDNA HS Assay kit (Thermo Fisher Scientific). Gel electrophoresis using Tapestation 2200 and D1000/High sensitivity D1000 screentapes (Agilent) was used to validate product size and purity of a subset of sequencing libraries. Sequencing libraries were prepared from the purified amplicon libraries using a second PCR. Each PCR reaction (25 mL) contained PCRBIO HiFi buffer (1x), PCRBIO HiFi Polymerase (1 U/reaction) (PCRBiosystems), adaptor mix (400 nM of each forward and reverse) and up to 10 ng of amplicon library template. PCR was conducted with the following program: initial denaturation at 95 °C for 2 min, 8 cycles of amplification (95 °C for 20 s, 55 °C for 30 s, 72 °C for 60 s) and a final elongation at 72 °C for 5 min. The resulting sequencing libraries were purified using the standard protocol for Agencourt Ampure XP Beads (Beckman Coulter) with a bead to sample ratio of 4:5. DNA was eluted in 25 mL of nuclease-free water (Qiagen). DNA concentration was measured using Qubit dsDNA HS Assay kit (Thermo Fisher Scientific). Gel electrophoresis using Tapestation 2200 and D1000/High sensitivity D1000 screentapes (Agilent) was used to validate product size and purity of a subset of sequencing libraries.

The purified sequencing libraries were pooled in equimolar concentrations and diluted to 2 nM. The samples were paired-end sequenced (2×300 bp) on a MiSeq (Illumina) using a MiSeq Reagent kit v3 (Illumina) following the standard guidelines for preparing and loading samples on the MiSeq. >10% PhiX control library was spiked in to overcome low complexity issues often observed with amplicon samples.

### Bioinformatic processing for fungal and bacterial gene regions

Forward and reverse reads were trimmed for quality using Trimmomatic v. 0.32^49^ with the settings SLIDINGWINDOW:5:3 and MINLEN: 225. The trimmed forward and reverse reads were merged using FLASH v. 1.2.7^50^ with the settings-m 10 −M 250. The trimmed reads were dereplicated and formatted for use in the UPARSE workflow^51^. The dereplicated reads were clustered, using the usearch v. 7.0.1090-cluster_otus command with default settings. Operational taxonomic unit (OTU) abundances were estimated using the usearch v. 7.0.1090-usearch_global command with-id 0.97 - maxaccepts 0 - maxrejects 0. Taxonomy was assigned using the RDP classifier^52^ as implemented in the parallel_assign_taxonomy_rdp.py script in QIIME^53^, using confidence 0.8 and the SILVA database, release 132^30^. The results were analysed in R v. 3.6.1^54^ through the Rstudio IDE using the ampvis package v.2.5.2^45^.

### Metagenome-assembled genomes – DNA extraction and sequencing

DNA was extracted using the DNeasy PowerSoil Pro Kit following the manufacturer’s recommendations (Qiagen, Germany), by DNASense ApS. A custom SPRi-bead size-selection protocol was further used to clean up and size-select (approx. cut-off 1500-2000 bp) DNA prior to Nanopore sequencing. Gel electrophoresis on Genomic DNA ScreenTapes and using the Tapestation 2200 (Agilent, USA) was used to validate DNA size distribution of a subset of DNA extracts. DNA concentration and purity were measured with the Qubit dsDNA HS Assay kit (Thermo Fisher Scientific, USA) and on the NanoDrop One (Thermo Fisher Scientific, USA).

The DNA was quantified using Qubit (Thermo Fisher Scientific, USA) and fragmented to approximately 550 bp using a Covaris M220 with microTUBE AFA Fiber screw tubes and the settings: Duty Factor 10%, Peak/Displayed Power 75W, Cycles/Burst 200, Duration 40 s and Temperature 20 °C. The fragmented DNA was used for metagenome preparation using the NEB Next Ultra II DNA library preparation kit. The DNA library was paired-end sequenced (2 x 151 bp) on a NovaSeq S4 system (Illumina, USA). A long-read sequencing library was prepared according to the SQK-LSK110 protocol (ONT, Oxford, United Kingdom). Approximately 50-100 fmole was loaded onto primed FLO-MIN106D (R9.4.1) flow cells and sequenced on the GridION platform using MinKNOW Release 22.05.7 (MinKNOW Core 5.1.0). Raw Illumina reads were filtered for PhiX using Usearch11^55^ and trimmed for adapters using Cutadapt v. 3.7^56^. Rasusa v. 0.6.1 was used to subset the Illumina data to 100 gb (gigabases)^57^. Raw Oxford Nanopore Fast5 files were basecalled in Guppy v. 6.1.5 using the Super-accurate basecalling algorithm. Basecalled fastq were was then adapter-trimmed in Porechop v.0.2.4 Porechop using default settings. NanoStat v.1.6.0^58^ was used to assess quality parameters of the basecalled data. The trimmed data were then filtered in Filtlong v. 0.2.0 with min_length set to 1000 bp and min_mean_q set to 95.

Draft metagenomes were assembled with Flye v.2.9^59^ by setting the metagenome parameter (– meta). The assembled metagenome was subsequently polished with quality-filtered ONT data using one round of polishing with Minimap2 v. 2.24^60^ and Racon v.1.4.20^61^ and two rounds of polishing with Medaka (v.1.6.1). The metagenome was finally polished with minimap2 v. 2.24^60^ and racon v.1.4.20^61^ using Illumina data. The metagenome was visualized in RStudio IDE (4.2.0 (2022-04-22)) running R version 4.0.4 (2021-02-15) and using the mmgenome2 R package v. 2.1.3. Metagenome-assembled genomes (MAGs) were manually extracted in mmgenome2 v. 2.2.1. Completeness and contamination values were assessed using CheckM v. 1.1.3^62^. MAG abundance was calculated using CoverM v. 0.6.1. MAGs were classified using the Genome Taxonomy Database toolkit v. 2.1.0 against the r207_v2 Genome Taxonomy Database^28, 63^. Ribosomale RNA genes were identified and extracted using Barrnap v. 0.9. MAGs were annotated using Prokka v. 1.14.6^64^.

### Analysis of Metagenome-assembled genomes

Putative resin acid-degrading proteins present in the MAGs were identified by using Basic Local Alignment Search Tool (BLAST) with protein sequences from *dit* cluster genes from *Pseudomonas abietaniphila* BKME-9 and *Paraburkholderia xenovorans* (Table S7) against the MAGs using an E-value of 1e^−10^ and sequence identity ≥30%. Putative carbohydrate active enzymes were identified using dbCAN^25^. Functional annotation of protein family databases (pfam) was performed in eggNOG-mapper using usegalaxy.eu^65, 66^.

### Isolation and identification of resin acid degrading bacteria

Resin acid degrading bacteria were isolated by serial dilution from one sample after the two weeks of bark degradation, that had been stored at −20 °C, using plates containing *Pseudomonas* Isolation Agar (PIA) (20.0 g/L beef extract peptone, 1.4 g/L magnesium chloride, 10 g/L potassium sulfate, 13.6 g/L agar, 20 mL/L glycerol). 5 g of the partially degraded bark was diluted in 45 mL of water, left to soak for 30 min at room temperature, and 100 mL inoculum from the resulting liquid was spread onto the plates with 10-fold dilutions up to 10^−5^. Plates were incubated at room temperature for five days. Obtained colonies were re-streaked on M9 minimal medium agar plates (15 g/L agar) containing 10 g/L of abietic acid. Single colonies were obtained by sequential streaking on fresh abietic acid plates, and for initial identification these were picked, boiled for 10 min, and used for PCR amplification of 16S rRNA and the signature *Pseudomonas* gene *rpoD*. Each PCR reaction (50 mL) contained Phusion polymerase (Thermo Scientific), 5X HF buffer (Thermo Scientific), water, 1 mL of boiled colonies, and 10 nM of each forward and reverse tailed primer T primers 27F (5′-AGAGTTTGATCMTGGCTCAG-3′) and 1470R (5′-TACGGYTACCTTGTTACGACT T-3′), and the partial *rpoD* sequence using primers PsEG30F (5’-ATYGAAATCGCCAARCG-3’) and PsEG790R (5’-CGGTTGATKTCCTTGA-3’), as previously described^67^. PCR was conducted with the following program: initial denaturation at 94 °C for 20 s, 35 cycles of amplification (94 °C for 20 s, 55 °C for 30 s, 72 °C for 60 s) and a final elongation at 72 °C for 7 min. The PCR products were analyzed by gel electrophoreses, purified, and sequenced using Sanger sequencing (Macrogen). Sequence results were identified using BLAST at NCBI^68^. An isolate, designated PIA16, was selected for genome sequencing and physiological tests, based on its ability to grow on abietic acid and sequence dissimilarity (16S rRNA and *rpoD*) to known species. The species was later named *Pseudomonas abieticivorans*, which is used henceforth.

### Genome sequencing of Pseudomonas abieticivorans

DNA extraction, sequencing, and analysis were performed by DNAsense ApS. DNA was extracted using the DNeasy PowerSoil Pro kit following the manufacturer’s recommendations (Qiagen) with minor modifications. Gel electrophoresis using Tapestation 2200 (Genomic DNA and D1000 screentapes, Agilent) was used to validate product size and purity of a subset of DNA extracts. DNA concentration was measured using the Qubit dsDNA HS Assay kit (Thermo Fisher Scientific). A rapid (SQK-RBK 110.96) library was prepared according to the manufacturer’s protocol, except for minor modifications (Oxford Nanopore Technologies). The library was loaded on a primed R9.4.1 MinION flow cell and sequenced on the GridION platform using MinKNOW Release 21.10.4. Sample DNA concentrations were measured using the Qubit dsDNA HS kit and the DNA quality was evaluated using TapeStation with the Genomic ScreenTape (Agilent Technologies). Sequencing libraries were prepared using the NEB Next Ultra II DNA library prep kit for Illumina (New England Biolabs) following the manufacturer’s protocol. Library concentrations were measured in triplicate using the Qubit dsDNA HS kit and library size estimated using TapeStation with D1000 HS Screen-Tape. The sequencing libraries were pooled in equimolar concentrations and diluted to 4 nM. The samples were paired end sequenced (2×301 bp) on a MiSeq (Illumina) using a MiSeq Reagent kit v3, 600 cycles (Illumina) following the standard guidelines for preparing and loading samples on the MiSeq. Fast5 raw data were basecalled and demultiplexed in Guppy (Oxford Nanopore Technologies) v. 6.0.1 using the sup algorithm. Basic read statistics including amount of data produced, median read quality and read N50 were assessed with NanoPlot v. 1.36.2^58^. Raw Illumina reads were filtered for PhiX using Usearch11^55^ and trimmed for adapters using cutadapt (v. 2.8^56^). A draft de novo assembly was produced with Flye v. 2.9-b1768^59^. The draft assembly was subsequently polished one time with Racon v. 1.4.20^61^ and Minimap2 v. 2.22-r1101^60^ and twice with Medaka v. 1.4.3 (Oxford Nanopore Technologies). The genome was finally polished with Minimap2 v. 2.22-r1101^60^ and Racon v.1.4.13^61^ using Illumina data. Assembly graph and draft assembly coverage was visualized and inspected with Bandage v. 0.8.1^69^. Genome annotation was conducted using Prokka (https://github.com/tseemann/prokka) v. 1.14.6^64^. Sample genomes were classified with the GTDB (Genome Taxonomy Database) tool Kit v. 1.7.0^70^ against the GTDB r202^63^. The genome was functionally annotated (COG) using WebMGA^71^.

### Phylogenetic and collinearity analysis

To compare the similarity between *P. abieticivorans* and known species, a phylogenetic analysis of all the *Pseudomonas* species reported in NCBI: GenBank^72^ was performed. BLASTn on NCBI web server^72^ was used to acquire 16S ribosomal RNA sequences, with *Pseudomonas* (taxid:286) sequences database selected, and limited to a maximum of 1000 target sequences. Multiple sequence alignment was executed using CLUSTALW with *msa* package^73^ in R v4.2.2^54^. Alignments were processed by using bios2msd^74^ and *adegenet*^75^ R packages. Phylogenetic trees were constructed using *ggtree*^76^ and *ggplot2*^77^ R packages. Tree estimation was calculated by the neighbour-joining method supported by ape^78^ package. Collinear analysis of *P. abieticivorans* compared to five species reported to be able to utilize resin acids (Table S11) was performed in MCScanX^79^. Protein-coding sequences were used. A local BLAST database was built for each species, to further run BLASTp^80^ with E-value less than 1e-10 and maximum 5 hits. In all the cases, *P. abieticivorans* was set as query. Default parameters were kept running MCScanX. Input files and output file analysis were carried out in R. Circos v0.69-9^81^ and used to visualize the collinear results.

### Morphology, physiological, and biochemical tests

Negative staining and transmission electron microscopy (TEM) were used to examine the morphology of *P. abieticivorans*. Formvar-coated copper grids (150 mesh) were rendered hydrophilic by glow discharge before applying 5 µL of cell suspension (OD_600_ = 0.5 in lysogeny broth; LB) for 3 min. The suspension was removed by blotting onto Whatman filter paper and the remaining bacteria were immediately fixed by applying 5 µL of 2% glutaraldehyde for 2 min. After blotting away the fixative and washing the grids twice with distilled water, the samples were stained for 1 min with 1% uranyl acetate. Excess stain was removed by blotting and the grids were air-dried before imaging. The samples were examined with a Talos L120C transmission electron microscope (Thermo Fisher Scientific) operating at 120 kV, and images were captured with a 4k × 4k BM-Ceta CMOS camera.

Growth at various temperatures (4, 20, 30, 37, 45 °C) was investigated on LB plates over 4 days and growth was assessed based on the presence of visible colonies. Growth based on salt concentration (NaCl 0-12% w/v) and the pH-growth range was assessed in LB medium by monitoring OD_600_ after 4 days of growth, with pH adjusted using NaOH or HCl. Fluorescence was studied by plating on King A or King B media (Millipore). Additional physiological tests were conducted using the API 20 NE and API ZYM systems according to the manufacturer’s instructions (bioMérieux). All tests were performed in duplicate.

### Chemotaxonomic analysis

Analysis of cellular fatty acids was carried out by DSMZ Services, Leibniz-Institut DSMZ – Deutsche Sammlung von Mikroorganismen und Zellkulturen GmbH (Braunschweig, Germany). The fatty acid methyl esters of the PIA16 strain were obtained from cells grown in tryptic soy broth (Millipore) at 28 °C for 24 h. Cellular fatty acids were analyzed after conversion into fatty acid methyl esters (FAMEs) by saponification, methylation, and extraction using minor modifications of established methods^82, 83^. The fatty acid methyl esters mixtures were analyzed by GC-FID using Sherlock Microbial Identification System (MIS) (MIDI, Microbial ID). Peaks were automatically integrated, and fatty acid names and percentages were calculated by the MIS Standard Software (Microbial ID).

### Growth on resin acids and resin acid analysis

Growth on dehydroabietic acid and abietic acid was done in cultures of M9 medium containing 100 mg/L of dehydroabietic acid (Megazyme ∼90%) or abietic acid (Sigma ∼75%) and inoculated with a starting optical density at 600 nm (OD_600_) of 0.01. The concentration of the resin acids (100 mg/L) was deliberately kept low to minimize background as resin acids are present as insoluble particles in the medium. Prior to inoculation, *P. abieticivorans* was grown in 10 mL LB medium at room temperature overnight. To follow growth and resin acid degradation over time, 1 mL of culture was sampled for evaluation of growth using protein concentration and degradation of resin acids using GC-MS. Intracellular protein concentration was used as a pseudo-value for growth due to the resin acids’ low solubility in water, making OD measurements unreliable. Briefly, cells were spun down, the pellet was resuspended in 1 M NaOH, boiled, and neutralized using HCl. Protein concentration was measured using the Bio-Rad BCA kit following the manufactures instruction. Resin acid extracts were analyzed using a modified method according to Bicho et al.^84^. Briefly, the resin acids were acidified using 40 µL HCl and spiked with an internal standard of 0.1 mg isopimaric acid (Carbosynth) and extracted three times using 1 mL ethyl acetate. Resin acids were dried over nitrogen and resuspended in 500 µL acetone and silylated using 25 µL BSTFA and TMCS (Sigma). Resin acid identification and quantification were performed by capillary GC (Agilent 7890A). Helium, the carrier gas, had a flow rate of 1 mL/min and the MS source was operated at 230 °C with the quadrupole at 150 °C. Analytes were separated using a HP-5 column with a temperature program starting at 70 °C, which was held for 2.25 min, was increased to 200 °C at 20 °C/min, thereafter 5 °C/min until the temperature reached 230 °C. The final ramp was at 35 °C min to 300 °C which was held for 10 min. Injector temperatures were at 300 °C. Helium, the carrier gas, had a flow rate of 1 mL/min and the MS source was operated at 230 °C with the quadrupole at 150 °C. The NIST MS Search Programme (Vers. 2.2) was used for identification using the library NIST/EPA/NIH Mass Spectral Library (NIST 11). Semi-quantification of the resin acids was done as using the internal standard and the following relationship: WS=AS×WI/AI, where W stands for the mass fraction, A for the area in the chromatogram, I the internal standard (isopimaric acid) and S the species, respectively.

The growth of *P. abieticivorans* on different types of resin acids was additionally analyzed using a Growth Profiler (Enzyscreen), using continuous imaging and pixel-based growth determination, in a 24-well plate (6×4) using 100 mg/L of each resin acid: dehydroabietic acid (Megazyme ∼90%), abietic acid (Sigma ∼75%), and isopimaric acid (Carbosynth). Growth was screened at 30 °C, and the cultures inoculated from an overnight culture of *P. abieticivorans* grown on LB broth to a starting OD_600_ of 0.01 in a total volume of 1.2 mL.

### Code availability

All the datasets and code used in this study for phylogenetics 16s rRNA and collinearity analysis are available at DOI:10.5281/zenodo.7596797.

### Data availability

All sequencing reads have been deposited at the National Center for Biotechnology Information (NCBI) under BioProject ID PRJNA912085. The 16S, ITS, and metagenomic reads have been deposited in the Sequence Read Archive (SRA) under the accession numbers SRR22745541-SRR22745556 (biosample accession SAMN32240041-SAMN32240056), SRR22745729-SRR22745736 (biosample accession SAMN32240041-SAMN32240056), and SRX18691017, SRX18691016 (biosample accession SAMN32218794) respectively (Table S13). The recovered MAGs have been deposited in GenBank under the accession numbers SAMN32218794 (biosample accession SAMN32233987-SAMN32234001) (Table S14). The *P. abieticivorans* PIA16 genome has been deposited in GenBank under the accession numbers GCA_023509015.1, ON945571.1 for the 16S rRNA sequence, and OP594298 for the *rpoD* sequence. *Pseudomonas abieticivorans* sp. nov. is deposited at the Culture collection University of Gothenburg (CCUG 76343T) and Deutsche Sammlung von Mikroorganismen und Zellkulturen GmbH (DSM 114633). Other data generated or analyzed during this study are included in the Supplementary Information files, and Source data is also provided.

## AUTHOR STATEMENTS

### Authors and contributors

Conceptualization, ASR, JL, PBP; Formal analysis, ASR, SV, AI, MH, ATR; Funding acquisition, ASR, MH, JL; Methodology, ASR, MH, PBP, JL; Project administration, ASR, JL; Resources, ASR, MH, JL; Supervision, JL, MH; Validation, ASR, ATR, MH, JL; Visualization, ASR, AI, ATR, SV, JL; Writing – original draft, ASR; Writing – review & editing, ASR, AI, ATR, EJK, MH, JL, PBP, SV.

### Conflicts of interest

The authors declare no conflict of interest.

## Funding information

Funding was provided by an energy-oriented basic research grant from the Swedish Energy Agency and the Swedish Research Council (project number 46559-1) and a research project grant from the Carl Trygger foundation (CTS 21:1424), awarded to JL, as well as two grants from the Adlerbertska Foundations, awarded to ASR and JL. EJK acknowledges funding from the Novo Nordisk Foundation (grant no. NNF20CC0035580).

## Acknowledgements

We would like to thank Dr. Marcel Taillefer for helpful discussions, and Holmen AB for providing spruce bark. We acknowledge the Centre for Cellular Imaging at the University of Gothenburg and the National Microscopy Infrastructure, NMI (VR-RFI 2019-00022) for assistance with electron microscopy.

## Abbreviations

ANI: average nucleotide identity
BLAST: basic local alignment search tool
Cazyme: carbohydrate-active enzyme
FID: flame ionization detector
GBDP: Genome Blast Distance Phylogeny
GC: gas chromatography
GH: glycoside hydrolase;
GHxx: glycoside hydrolase family xx
GT: glycosyltransferase
GTDB: Genome Taxonomy Database
HPAEC-PAD: high-performance anion-exchange chromatography with pulsed amperometric detection
LB: Lysogeny broth
MAG: metagenome-assembled genome
PIA: *Pseudomonas* isolation agar
PUL: polysaccharide utilization locus
TYGS: Type (Strain) Genome Server

## Notes

### Competing Interest Statement

The authors have declared no competing interest.

