## Supplemental file for "Resin acids play key roles in shaping microbial communities during degradation of spruce bark"

**Affiliations:**

**INDEX**

### Supplementary tables

**Table S1.** Intervals of integration for the main classes of extractive compounds and internal standards in the spruce bark extractives using GC-MS and the corresponding column used.

| **Class of compounds** | | | |
|---|---|---|---|
| Name | Acronym | Retention time interval (min) | Column (length) |
| Fatty acids | FA | 14.5-16.9 | HP-5ms (30 m) |
| Resin acids | RA | 17.0-18.0 | HP-5ms (30 m) |
| Sterols | ST | 24.1-25.6 | HP-5ms (30 m) |
| Steryl esters | SE | 17.3-18.9 | HP-1 SIM/Dist(5 m) |
| Triglycerides | TG | 19.7-22.0 | HP-1 SIM/Dist (5 m) |
| **Internal standards** | | | |
| Name | Acronym | Retention time interval (min) | Column (length) |
| Heptadecaonic acid | INS1 | 20.05 – 20.14 | HP-5ms (30 m) |
| Cholesterylheptadecanoate | INS2 | 16.9-17.4 | HP-1 SIM/Dist(5 m) |
| 1.3-dipalmitoyl-2 oleyl glycerol | INS3 | 18.9-19.7 | HP-1 SIM/Dist(5 m) |

**Table S2**. Identification of individual compounds in spruce bark extract by GC-MS/FID on a HP-5ms column after derivatization with BSTFA/TMCS/pyridine in the biotic sample at two weeks growth, and the abiotic sample at week zero.

| Retention time (min) | Compound name | CAS nr | Formula | Match factor | Reverse match | Probability (%) |
|---|---|---|---|---|---|---|
| 14.2 | Hexadecanoic acid | 57-10-3 | C16H32O2 | 789 | 826 | 65 |
| 14.85 | Heptadecanoic acid | 506-12-7 | C17H34O2 | 705 | 854 | 57.5 |
| 15.54 | 9,12,15-Octadecatrienoic acid | 1955-33-5 | C18H30O2 (C18:3) | 803 | 843 | 20.8 |
| 15.702 | 9,12-Octadecadienoic acid | 2197-37-7 | C18H34O2 (C18:2) | 882 | 947 | 63.4 |
| 15.745 | (9,Z)-Octadec-9-enoic acid | 112-80-1 | C18H34O2 (C18:1) | 897 | 943 | 22.8 |
| 15.955 | Stearic acid | | C18 | 497 | 705 | 24.7 |
| 16.655 | Pimaric acid | 127-27-5 | C20H30O2 | 861 | 887 | 72.8 |
| 16.785 | Isopimaric acid | 5835-26-7 | C20H30O2 | 868 | 911 | 66.6 |
| 17.075 | Dehydroabietic acid | 1740-19-8 | C20H28O2 | 816 | 835 | 71.9 |
| 17.54 | Abietic acid | 514-10-3 | C20H30O2 | 868 | 889 | 82.3 |
| 19.098 | 7-oxodehydroabietic acid | 18684-55-4 | C20H26O3 | 783 | 836 | 91.7 |

*900 or greater is considered an excellent match; 800–900 a good match; 700–800 a fair match and less than 600 is a very poor match

**Table S3. Sequencing statistics (fungi).** Extraction conc. is the concentration of the extracted DNA (ng/µL). Library conc. is the concentration of the sequencing library (ng/µL). Reads is the number of DNA reads after sequencing, quality control (QC) and bioinformatic processing. Observed amplicon sequence variants (ASVs) is the number of observed, unique ASVs in each sample.

| **Sample name** | **Week** | **Extraction concentration (ng/uL)** | **Amplication conc (ng/uL)** | **Library conc (ng/uL)** | **Reads** | **Observed ASVs** |
|---|---|---|---|---|---|---|
| w0-s3 | 0 | 6.3 | 7.9 | 10.6 | 20978 | 328 |
| w0-s1 | 0 | 7.3 | 10.4 | 4.2 | 21679 | 329 |
| w2-s6 | 2 | 96.3 | 13.2 | 13.8 | 19329 | 281 |
| w2-s5 | 2 | 61.6 | 12.7 | 8.5 | 22553 | 276 |
| w4-s9 | 4 | 65.9 | 15.4 | 13.1 | 25210 | 269 |
| w4-s8 | 4 | 62.4 | 11.9 | 9 | 28105 | 245 |
| w8-s11 | 8 | 68.6 | 10.8 | 7 | 19047 | 236 |
| w8-s12 | 8 | 61.4 | 9.4 | 3.7 | 23858 | 229 |
| w12-s15 | 12 | 58 | 7 | 10.2 | 25422 | 238 |
| w12-s14 | 12 | 53 | 7.3 | 11 | 25795 | 225 |
| w16-s18 | 16 | 38.6 | 9.9 | 12.5 | 20731 | 218 |
| w16-s17 | 16 | 67.4 | 8.5 | 9.5 | 21673 | 226 |
| w20-s20 | 20 | 45.5 | 6.5 | 7.4 | 25009 | 220 |
| w20-s21 | 20 | 58.7 | 13.2 | 6.9 | 29052 | 240 |
| w24-s23 | 24 | 55 | 3.9 | 9.3 | 22479 | 223 |
| w24-s24 | 24 | 52.5 | 7.8 | 8.5 | 26682 | 221 |

**Table S4. Sequencing statistics (bacteria).** Extraction conc. is the concentration of the extracted DNA (ng/µL). Library conc. is the concentration of the sequencing library (ng/µL). Reads is the number of DNA reads after sequencing, QC and bioinformatic processing. Observed OTUs is the number of observed, unique OTUs in each sample.

| **Sample name** | **Week** | **Extraction concentration (ng/uL)** | **Amplication conc (ng/uL)** | **Library conc (ng/uL)** | **Reads** | **Observed OTUs** |
|---|---|---|---|---|---|---|
| w0-s3 | 0 | 6.3 | 7.9 | 10.6 | 38037 | 377 |
| w0-s1 | 0 | 7.3 | 10.4 | 4.2 | 46811 | 457 |
| w2-s5 | 2 | 96.3 | 13.2 | 13.8 | 38986 | 283 |
| w2-s6 | 2 | 61.6 | 12.7 | 8.5 | 51433 | 292 |
| w4-s9 | 4 | 65.9 | 15.4 | 13.1 | 38597 | 317 |
| w4-s8 | 4 | 62.4 | 11.9 | 9 | 54529 | 297 |
| w8-s11 | 8 | 68.6 | 10.8 | 7 | 57777 | 325 |
| w8-s12 | 8 | 61.4 | 9.4 | 3.7 | 61646 | 349 |
| w12-s15 | 12 | 58 | 7 | 10.2 | 48408 | 356 |
| w12-s14 | 12 | 53 | 7.3 | 11 | 49855 | 356 |
| w16-s18 | 16 | 38.6 | 9.9 | 12.5 | 51583 | 354 |
| w16-s17 | 16 | 67.4 | 8.5 | 9.5 | 62390 | 363 |
| w20-s21 | 20 | 45.5 | 6.5 | 7.4 | 64313 | 371 |
| w20-s20 | 20 | 58.7 | 13.2 | 6.9 | 70628 | 366 |
| w24-s24 | 24 | 55 | 3.9 | 9.3 | 59273 | 358 |
| w24-s23 | 24 | 52.5 | 7.8 | 8.5 | 67774 | 363 |

**Table S5. Bark two-week metagenome assembly statistics**. The number of contigs denotes the number of de novo assembled contigs, with length total (Mb) denoting the total length (in megabases). Length maximum (Mb), length mean (bp) and length minimum (bp) denote the maximum, mean and minimum contig lengths (in basepairs). N50 (bp) is defined as the sequence length of the shortest contig at 50% of the total assembly length. Mean GC content (%) denotes the mean content of the G- and C-nucleotides in the assembly.

| Summary | Bark assembly |
|---|---|
| Number of contigs | 15,474 |
| Total length (Mb) | 430 |
| Length maximum (Mb) | 4.32 |
| Length mean (bp) | 27,784 |
| Length minimum (bp) | 344 |
| N50 (bp) | 49,770 |
| Mean GC content (%) | 59 |
| Data (Gbp) - Oxford Nanopore | 16 |
| Reads (mio.) - Oxford Nanopore | 2.9 |
| Q-score - Oxford Nanopore | 14.5 |
| Data (Gbp) - Illumina | 340 |
| Reads (mio.) - Illumina | 2,266 |
| Q-score - Illumina | 33.8 |

**Table S6. MAG statistics.** MAG ID and MAG size refer to the bin identification number and the size of the MAG in megabases, respectively. Contigs denotes the number of contiguous DNA elements associated with each MAG. Compl. (%) is the estimated genome completeness based on the presence/absence of essential lineage-specific marker genes. Cont. (%) is the estimated contamination based on the presence of multiple single-copy marker genes. N50 (mb) is defined as the sequence length (megabases) of the shortest contig at 50% of the total MAG length. MAG quality refers to the quality of the extracted genome based on the MIMAG standard (Minimal Information about a Metagenome-assembled genome), with HQ denoting high-quality (>90% completion, <5% contamination, presence of 5S, 16S and 23S rRNA and a minimum of 18 tRNA detected) MAGs. GTDB taxonomy refers to the GTDB (Genome Taxonomy Database) classification at the highest taxonomic resolution assigned to a specific MAG.

| MAG ID | MAG size (mbp) | contigs | N50 (mbp) | Compl (%) | Cont (%) | GTDB taxonomy | MAG quality |
|---|---|---|---|---|---|---|---|
| Bin1 | 7.16 | 5 | 2.21 | 99.34 | 0.15 | *Nitrospirillum* | HQ |
| Bin2 | 4.32 | 1 | 4.32 | 98.45 | 2.43 | *Pseudoxanthomonas*_A | HQ |
| Bin3 | 2.56 | 9 | 1.42 | 98.44 | 1.69 | GAS474 | HQ |
| Bin4 | 3.24 | 9 | 2.08 | 99.35 | 0.10 | JAATFR01 | HQ |
| Bin5 | 3.37 | 10 | 0.38 | 95.66 | 8.41 | UBA2020 | MQ |
| Bin6 | 8.09 | 8 | 1.40 | 96.51 | 1.56 | *Paraburkholderia tropica* | HQ |
| Bin7 | 7.06 | 21 | 0.45 | 97.47 | 1.69 | *Paraburkholderia* | HQ |
| Bin8 | 4.09 | 4 | 1.10 | 99.61 | 0.00 | *Neorhizobium* | HQ |
| Bin9 | 6.24 | 34 | 0.34 | 98.39 | 1.23 | *Pantoea* | HQ |
| Bin10 | 4.27 | 5 | 3.59 | 100.00 | 0.86 | Tous-C9LFEB | HQ |
| Bin11 | 4.79 | 10 | 0.56 | 96.55 | 0.86 | *Edaphobacter* | HQ |
| Bin12 | 3.01 | 31 | 0.19 | 90.52 | 2.76 | Tous-C9LFEB | MQ |
| Bin13 | 3.59 | 35 | 0.13 | 97.78 | 1.72 | *Arachidicoccus* | HQ |
| Bin14 | 6.45 | 16 | 0.61 | 99.18 | 1.19 | *Pseudomonas*_E sp002080045 | HQ |
| Bin15 | 6.62 | 69 | 0.16 | 90.67 | 3.08 | *Pseudomonas*_E | HQ |

**Table S7.** NCBI accession number of the genes from the *dit* cluster genes from *Pseudomonas abietaniphila* BKME-9 and *Paraburkholderia xenovorans* LB400.

| Gene | NBCI accession number | Description | Microorganism |
|---|---|---|---|
| *ditI* | AAD21071.1 | Dehydrogenase/reductase | *Pseudomonas abietaniphila* BKME-9[1] |
| *ditA2* | AAD21061.1 | β Subunit of the ring-hydroxylating dioxygenase | *Pseudomonas abietaniphila* BKME-9[1] |
| *ditA1* | AAD21063.1 | α Subunit of the ring-hydroxylating dioxygenase | *Pseudomonas abietaniphila* BKME-9[1] |
| *ditH* | AAD21070.1 | Isomerase/decarboxylase | *Pseudomonas abietaniphila* BKME-9[1] |
| *ditG* | AAD21069.1 | Dehydrogenase/reductase | *Pseudomonas abietaniphila* BKME-9[1] |
| *ditF* | AAD21068.1 | Sterol carrier-like protein | *Pseudomonas abietaniphila* BKME-9[1] |
| *ditR* | AAD21072.1 | IclR-type transcription regulator | *Pseudomonas abietaniphila* BKME-9[1] |
| *ditE* | AAD21067.1 | Permease of the major facilitator superfamily | *Pseudomonas abietaniphila* BKME-9[1] |
| *ditD* | AAD21066.1 | Isomerase/decarboxylase | *Pseudomonas abietaniphila* BKME-9[1] |
| *ditC* | AAD21065.1 | Extradiol cleavage dioxygenase | *Pseudomonas abietaniphila* BKME-9[1] |
| *ditB* | AAD21064.1 | Dehydrogenase/reductase | *Pseudomonas abietaniphila* BKME-9[1] |
| *ORF2* | AAD21074.1 | | *Pseudomonas abietaniphila* BKME-9[1] |
| *ditA3* | AAD21062.1 | Ferredoxin component of ring-hydroxylating dioxygenase | *Pseudomonas abietaniphila* BKME-9[1] |
| *ORF3* | AAR83736.1 | | *Pseudomonas abietaniphila* BKME-9[2] |
| *ORF4* | AAR83737.1 | | *Pseudomonas abietaniphila* BKME-9[2] |
| *ditJ* | AAD21073.2 | CoA ligase | *Pseudomonas abietaniphila* BKME-9[2] |
| *ditK* | AAR83744.1 | Transcriptional regulator, TetR family | *Pseudomonas abietaniphila* BKME-9[2] |
| *ditL* | AAR83743.1 | Hypothetical protein | *Pseudomonas abietaniphila* BKME-9[2] |
| *ditM* | AAR83742.1 | Hydrolase | *Pseudomonas abietaniphila* BKME-9[2] |
| *ditN* | AAR83741.1 | 3-hydroxyacyl CoA dehydrogenase | *Pseudomonas abietaniphila* BKME-9 |
| *ditO* | AAR83740.1 | Thiolase | *Pseudomonas abietaniphila* BKME-9[2] |
| *ditP* | AAR83739.1 | Conserved hypothetical protein | *Pseudomonas abietaniphila* BKME-9[2] |
| *ditQ* | AAR83738.1 | Cytochrome P450 | *Pseudomonas abietaniphila* BKME-9[2] |
| *ditA2* | ABE36485.1 | β Subunit of the ring-hydroxylating dioxygenase | *Paraburkholderia xenovorans* LB400[3] |
| *ditA1* | ABE36484.1 | α Subunit of the ring-hydroxylating dioxygenase | *Paraburkholderia xenovorans* LB400[3] |
| *ditH* | ABE36483.1 | Isomerase/decarboxylase | *Paraburkholderia xenovorans* LB400[3] |
| *ditG* | ABE36481.1 | Dehydrogenase/reductase | *Paraburkholderia xenovorans* LB400[3] |

| | | | |
|---|---|---|---|
| *ditF* | ABE36479.1 | Sterol carrier-like protein | *Paraburkholderia xenovorans* LB400[3] |
| *ditR* | ABE36505.1 | IclR-type transcription regulator | *Paraburkholderia xenovorans* LB400[3] |
| *ditD* | ABE36506.1 | Isomerase/decarboxylase | *Paraburkholderia xenovorans* LB400[3] |
| *ditC* | ABE36538.1 | Extradiol cleavage dioxygenase | *Paraburkholderia xenovorans* LB400[3] |
| *ditB* | ABE36537.1 | Dehydrogenase/reductase | *Paraburkholderia xenovorans* LB400[3] |
| *ditA3* | ABE36536.1 | Ferredoxin component of ring-hydroxylating dioxygenase | *Paraburkholderia xenovorans* LB400[3] |
| *ditK* | ABE36491.1 | Transcriptional regulator, TetR family | *Paraburkholderia xenovorans* LB400[3] |
| *ditM* | ABE36493.1 | Hydrolase | *Paraburkholderia xenovorans* LB400[3] |
| *ditN* | ABE36494.1 | 3-hydroxyacyl CoA dehydrogenase | *Paraburkholderia xenovorans* LB400[3] |
| *ditO* | ABE36495.1 | Thiolase | *Paraburkholderia xenovorans* LB400[3] |
| *ditP* | ABE36496.1 | Conserved hypothetical protein | *Paraburkholderia xenovorans* LB400[3] |
| *ditQ* | ABE36497.1 | Cytochrome P450 | *Paraburkholderia xenovorans* LB400[3] |
| *ditU* | ABE36529.1 | Cytochrome P450 | *Paraburkholderia xenovorans* LB400[3] |

**Table S8**.Genomic characteristics of *P. abieticivorans*.

| **Attribute** | **Genome** |
|---|---|
| Genbank ID | GCA_023509015.1 |
| Size (bp) | 6,715,763 |
| No. scaffolds/contigs | 1 |
| Cov. (fold) | 149 |
| G+C content (mol%) | 62 |
| Total genes | 6,087 |
| Protein-coding genes | 5,987 |
| KO numbers | 3,449 |
| tRNA | 74 |
| rRNA (5S, 16S, 23S) | 8, 7, 7 |
| Pseudogenes[a] | 139 |

[a]The number of total pseudogenes indicated includes genes with ambiguous residues, frameshifted genes, incomplete genes, genes with internal stops or other multiple problems.

**Table S9**. Number of genes associated with the general COG functional categories.

| Code | Value | % of total | Description |
|---|---|---|---|
| J | 245 | 4.09 | Translation |
| A | 25 | 0.4 | RNA processing and modification |
| K | 231 | 3.9 | Transcription |
| L | 238 | 3.97 | Replication, recombination and repair |
| B | 19 | 0.3 | Chromatin structure and dynamics |
| D | 72 | 1.2 | Cell cycle control, mitosis and meiosis |
| Y | 2 | 0.03 | Nuclear structure |
| V | 46 | 0.8 | Defense mechanism |
| T | 152 | 2.5 | Signal transduction mechanisms |
| M | 188 | 3.2 | Cell wall/membrane biogenesis |
| N | 96 | 1.6 | Cell motility |
| Z | 12 | 0.2 | Cytoskeleton |
| W | 1 | 0.02 | Extracellular structures |
| U | 158 | 2.7 | Intracellular trafficking and secretion |
| O | 203 | 3.4 | Posttranslational modification, protein turnover, chaperones |
| C | 258 | 4.3 | Energy production and conversion |
| G | 230 | 3.8 | Carbohydrate transport and metabolism |
| E | 270 | 4.5 | Amino acid transport and metabolism |
| F | 95 | 1.6 | Nucleotide transport and metabolism |
| H | 179 | 2.9 | Coenzyme transport and metabolism |
| I | 94 | 1.57 | Lipid transport and metabolism |
| P | 212 | 3.5 | Inorganic transport and metabolism |
| Q | 88 | 1.47 | Secondary metabolites biosynthesis, transport and catabolism |
| R | 702 | 11.8 | General function prediction only |
| S | 1347 | 22.46 | Function unknown |

**Table S10.** Phenotypic characteristics distinguishing *P. abieticivorans* from phylogenetically closely related *Pseudomonas* type strains. *P. abieticivorans* PIA16$^T$ (1), *P. baetica* a390$^T$(2), *P. putida* ICMP/NBRC 14164$^T$ (3), *P. koreensis* Ps 9-14$^T$ (4), *P. reinekei* MT1$^T$ (5). (data from;Lopez 2012[4]; Palleroni, 2005 [5];Kwon 2003[6]; Camara 2007[7]) All data are from this study unless indicated otherwise., All strains were grown on trypticase soy broth agar prior to fatty acid analysis. +, positive, -, negative; w, weak; tr, trace (<1%).

| **Characteristics** | **1** | **2** | **3** | **4** | **5** |
|---|---|---|---|---|---|
| Temperature | 4-30 | 4-30 | 4-30 | 4-30 | 4-30 |
| Hydrolysis (β-glucosidase) (esculin) | W | - | - | - | - |
| Hydrolysis (protease) (gelatin) | - | + | - | + | - |
| Assimilation (arabinose) | + | + | - | + | + |
| Assimilation (mannose) | - | + | - | + | - |
| Assimilation (*N*-acetylglucosamine) | - | + | - | + | - |
| Assimilation (phenylacetic acid) | - | - | + | - | + |
| **Enzyme** | | | | | |
| Alkaline phosphatase | + | + | - | - | w |
| Valine arylamidase | + | - | - | - | - |
| Naphthol-AS-BI-phosphohydrolase | + | - | + | + | + |
| Catalase | - | + | - | + | - |
| **Cellular fatty acid composition** | | | | | |
| $C_{12:0}$ | 4.11 | 1.68 | 2.6 | 3.9 | 3.55 |
| $C_{14:0}$ | tr | tr | tr | tr | tr |
| $C_{16:0}$ | 31.57 | 29.43 | 30.5 | 33.5 | 39.48 |
| $C_{10:0}$ 3-OH | 3 | 3.44 | 4 | 4.2 | 3.34 |
| $C_{12:0}$ 2-OH | 3.05 | 5.54 | 4.9 | 3.5 | 4.28 |
| $C_{12:0}$ 3-OH | 3.49 | 3.23 | 3.9 | 3.8 | 4.78 |
| $C_{17:0}$ cyclo | 12.79 | 3.15 | 12.3 | 4.6 | 22.25 |
| Summed feature 3:$C_{16:1}$ ω7c/ω6c | 26.12 | 39.50 | 23.7 | 35.3 | 12.22 |
| Summed feature 8: $C_{18:1}$ω7c/ω6c | 13.96 | 12.55 | 15.2 | 11.5 | 10.35 |
| DNA G+C content (mol%) | 61.5 | 58.5 | 62 | 60 | 59 |
| 16S rRNA sequence similarity to strain PIA16 | 100 | 98.66 | 98.61 | 98.42 | 98.39 |
| *rpoD* sequence similarity to PIA16 | 100 | 84.31 | 88.38 | 88.68 | 88.91 |
| ANI similarity to strain PIA16 | 100 | 81.4 | 81.6 | 81.6 | 81.5 |

**Table S11.** *Pseudomonas sp.* used for collinear analysis including strain, taxonomic identification, and NCBI assembly ID.

| id | Name | Strain | Taxid | NCBI assembly id |
|---|---|---|---|---|
| pp1 | *Pseudomonas abieticivorans* | PIA16 | 2931382 | GCF_023509015.1 |
| pr1 | *Pseudomonas resinovorans* | NBRC 106553 | 1245471 | GCF_000412695.1 |
| pm1 | *Pseudomonas multiresinivorans* | | 95301 | GCF_012971725.1 |
| pv1 | *Pseudomonas vancouverensis* | LMG 20222 | 95300 | GCF_900105825.1 |
| pa1-54 | *Pseudomonas abietaniphila* | ATCC 700689 | 89065 | GCF_900100795.1 |
| px1-3 | *Paraburkholderia xenovorans* | LB400 | 266265 | GCF_000013645.1 |

**Table S12. Blast of *dit* cluster-encoded proteins against *Pseudomonas abieticivorans*.** Hit denotes the name of the sequence found in the BLAST search and description is the Prokka-annotated description of said identified sequence. E-value is the measure of quality of the match. Higher E-values indicate that BLAST found a less homologous sequence. Identity % denotes the percentage of identical residues in the query and hit sequence.

| Gene | Query NCBI accession number | PIA16 NCBI accession number | Description | E-value | Percent identity % |
|---|---|---|---|---|---|
| *ditI* | AAD21071.1 | WP_249674695.1 | Diacetyl reductase [(S)-acetoin | 6.042e-152 | 83.83 |
| *ditA2* | AAD21061.1 | WP_249674693.1 | Biphenyl dioxygenase subunit | 5.864e-121 | 86.70 |
| *ditA1* | AAD21063.1 | WP_249674699.1 | Biphenyl 2,3-dioxygenase sub | 0 | 93.09 |
| *ditH* | AAD21070.1 | WP_249674700.1 | putative protein YisK | 0 | 82.57 |
| *ditG* | AAD21069.1 | WP_249674701.1 | 3-oxoacyl-[acyl-carrier-protein | 2.814e-105 | 71.74 |
| *ditF* | AAD21068.1 | WP_249674704.1 | hypothetical protein | 0 | 88.06 |
| *ditR* | AAD21072.1 | WP_249674706.1 | Bacterial transcriptional regulator | 8.720e-139 | 72.59 |
| *ditE* | AAD21067.1 | WP_249674707.1 | Enterobactin exporter EntS | 1.388e-148 | 63.08 |
| *ditD* | AAD21066.1 | WP_249674708.1 | putative protein | 2.649e-144 | 68.71 |
| *ditC* | AAD21065.1 | WP_249674709.1 | Iron-dependent extradiol | 0 | 82.32 |
| *ditB* | AAD21064.1 | WP_249674710.1 | Cyclopentanol dehydrogenase | 1.769e-140 | 81.27 |
| *ditA3* | AAD21062.1 | WP_249674711.1 | 4Fe-4S single cluster domain | 2.969e-17 | 81.33 |
| ORF2 | AAD21074.1 | WP_249674712.1 | Long-chain-fatty-acid--CoA | 0 | 76.29 |
| ORF3 | AAR83736.1 | WP_249674685.1 | Crotonobetainyl-CoA reductase | 0 | 78.53 |
| ORF4 | AAR83737.1 | WP_249674686.1 | Amidohydrolase | 0 | 90.11 |
| *ditQ* | AAR83738.1 | WP_249674687.1 | Putative cytochrome P450 | 0 | 84.47 |
| *ditP* | AAR83739.1 | WP_249674688.1 | hypothetical protein | 7.441e-69 | 66.87 |
| *ditO* | AAR83740.1 | WP_249674689.1 | Beta-ketoadipyl-CoA thiolase | 0 | 78.77 |
| *ditN* | AAR83741.1 | WP_249674690.1 | 3-hydroxybutyryl-CoA | 3.404e-180 | 80.13 |
| *ditM* | AAR83742.1 | WP_249674691.1 | Fumarylacetoacetate (FAA) | 9.019e-179 | 83.04 |
| *ditL* | AAR83743.1 | WP_249674692.1 | 2-keto-4-carboxy-3-hexenedioate | 0 | 85.80 |
| *ditK* | AAR83744.1 | WP_249674693.1 | hypothetical protein | 3.843e-127 | 84.95 |
| *ditU* | ABE36529.1 | WP_249674723.1 | Putative cytochrome P450 | 0 | 64.30 |

**Table S13.** Description of Sequence Read Archive (SRA) accession numbers of each sample used for amplicon sequencing.

| SeqID | Libtype | Sample name | Week | Accession number |
|---|---|---|---|---|
| MQ201204-183 | ITS2 | w0-sample1 | 0 | https://www.ncbi.nlm.nih.gov/sra/SRX18706875 |
| MQ201204-184 | ITS2 | w0-sample3 | 0 | https://www.ncbi.nlm.nih.gov/sra/SRX18706876 |
| MQ201204-185 | ITS2 | w2-sample5 | 2 | https://www.ncbi.nlm.nih.gov/sra/SRX18706883 |
| MQ201204-186 | ITS2 | w2-s6 | 2 | https://www.ncbi.nlm.nih.gov/sra/SRX18706884 |
| MQ201204-187 | ITS2 | w4-s8 | 4 | https://www.ncbi.nlm.nih.gov/sra/SRX18706885 |
| MQ201204-188 | ITS2 | w4-s9 | 4 | https://www.ncbi.nlm.nih.gov/sra/SRX18706886 |
| MQ201204-189 | ITS2 | w8-s11 | 8 | https://www.ncbi.nlm.nih.gov/sra/SRX18706887 |
| MQ201204-190 | ITS2 | w8-s12 | 8 | https://www.ncbi.nlm.nih.gov/sra/SRX18706888 |
| MQ201204-191 | ITS2 | w12-s14 | 12 | https://www.ncbi.nlm.nih.gov/sra/SRX18706889 |
| MQ201204-192 | ITS2 | w12-s15 | 12 | https://www.ncbi.nlm.nih.gov/sra/SRX18706890 |
| MQ201204-193 | ITS2 | w16-s17 | 16 | https://www.ncbi.nlm.nih.gov/sra/SRX18706877 |
| MQ201204-194 | ITS2 | w16-s18 | 16 | https://www.ncbi.nlm.nih.gov/sra/SRX18706878 |
| MQ201204-195 | ITS2 | w20-s20 | 20 | https://www.ncbi.nlm.nih.gov/sra/SRX18706879 |
| MQ201204-196 | ITS2 | w20-s21 | 20 | https://www.ncbi.nlm.nih.gov/sra/SRX18706880 |
| MQ201204-197 | ITS2 | w24-s23 | 24 | https://www.ncbi.nlm.nih.gov/sra/SRX18706881 |
| MQ201204-198 | ITS2 | w24-s24 | 24 | https://www.ncbi.nlm.nih.gov/sra/SRX18706882 |
| MQ201204-213 | abV4-C | w0-sample1 | 0 | https://www.ncbi.nlm.nih.gov/sra/SRX18706695 |
| MQ201204-214 | abV4-C | w0-sample3 | 0 | https://www.ncbi.nlm.nih.gov/sra/SRX18706696 |
| MQ201204-215 | abV4-C | w2-sample5 | 2 | https://www.ncbi.nlm.nih.gov/sra/SRX18706703 |
| MQ201204-216 | abV4-C | w2-s6 | 2 | https://www.ncbi.nlm.nih.gov/sra/SRX18706704 |
| MQ201204-217 | abV4-C | w4-s8 | 4 | https://www.ncbi.nlm.nih.gov/sra/SRX18706705 |
| MQ201204-218 | abV4-C | w4-s9 | 4 | https://www.ncbi.nlm.nih.gov/sra/SRX18706706 |
| MQ201204-219 | abV4-C | w8-s11 | 8 | https://www.ncbi.nlm.nih.gov/sra/SRX18706707 |
| MQ201204-220 | abV4-C | w8-s12 | 8 | https://www.ncbi.nlm.nih.gov/sra/SRX18706708 |
| MQ201204-221 | abV4-C | w12-s14 | 12 | https://www.ncbi.nlm.nih.gov/sra/SRX18706709 |
| MQ201204-222 | abV4-C | w12-s15 | 12 | https://www.ncbi.nlm.nih.gov/sra/SRX18706710 |
| MQ201204-223 | abV4-C | w16-s17 | 16 | https://www.ncbi.nlm.nih.gov/sra/SRX18706697 |
| MQ201204-224 | abV4-C | w16-s18 | 16 | https://www.ncbi.nlm.nih.gov/sra/SRX18706698 |
| MQ201204-225 | abV4-C | w20-s20 | 20 | https://www.ncbi.nlm.nih.gov/sra/SRX18706699 |
| MQ201204-226 | abV4-C | w20-s21 | 20 | https://www.ncbi.nlm.nih.gov/sra/SRX18706700 |
| MQ201204-227 | abV4-C | w24-s23 | 24 | https://www.ncbi.nlm.nih.gov/sra/SRX18706701 |
| MQ201204-228 | abV4-C | w24-s24 | 24 | https://www.ncbi.nlm.nih.gov/sra/SRX18706702 |

**Table S14**. Metagenome-assembled genomes (MAGs) NCBI accession numbers from https://www.ncbi.nlm.nih.gov/sra/SRX18691017 and https://www.ncbi.nlm.nih.gov/sra/SRX18691016.

| MAG ID | Biosample ID | GenBank ID | Link |
|---|---|---|---|
| Bin1 | SAMN32241433 | GCA_028283825.1 | https://www.ncbi.nlm.nih.gov/nuccore/JAPWJW000000000 |
| Bin2 | SAMN32241434 | GCA_027498035.1 | https://www.ncbi.nlm.nih.gov/nuccore/2418877976 |
| Bin3 | SAMN32241435 | GCA_028283855.1 | https://www.ncbi.nlm.nih.gov/nuccore/JAPWJX000000000 |
| Bin4 | SAMN32241436 | GCA_028283765.1 | https://www.ncbi.nlm.nih.gov/nuccore/JAPWJY000000000 |
| Bin5 | SAMN32241437 | GCA_028283745.1 | https://www.ncbi.nlm.nih.gov/nuccore/JAPWJZ000000000 |
| Bin6 | SAMN32241438 | GCA_028283795.1 | https://www.ncbi.nlm.nih.gov/nuccore/JAPWKA000000000 |
| Bin7 | SAMN32241439 | GCA_028283785.1 | https://www.ncbi.nlm.nih.gov/nuccore/JAPWKB000000000 |
| Bin8 | SAMN32241440 | GCA_028283725.1 | https://www.ncbi.nlm.nih.gov/nuccore/JAPWKC000000000 |
| Bin9 | SAMN32241441 | GCA_028283705.1 | https://www.ncbi.nlm.nih.gov/nuccore/JAPWKD000000000 |
| Bin10 | SAMN32241442 | GCA_028283655.1 | https://www.ncbi.nlm.nih.gov/nuccore/JAPWKE000000000 |
| Bin11 | SAMN32241443 | GCA_028283685.1 | https://www.ncbi.nlm.nih.gov/nuccore/JAPWKF000000000 |
| Bin12 | SAMN32241444 | GCA_028283625.1 | https://www.ncbi.nlm.nih.gov/nuccore/JAPWKG000000000 |
| Bin13 | SAMN32241445 | GCA_028283605.1 | https://www.ncbi.nlm.nih.gov/nuccore/JAPWKH000000000 |
| Bin14 | SAMN32241446 | GCA_028283645.1 | https://www.ncbi.nlm.nih.gov/nuccore/JAPWKI000000000 |
| Bin15 | SAMN32241447 | GCA_028283575.1 | https://www.ncbi.nlm.nih.gov/nuccore/JAPWKJ000000000 |

### Supplementary figures

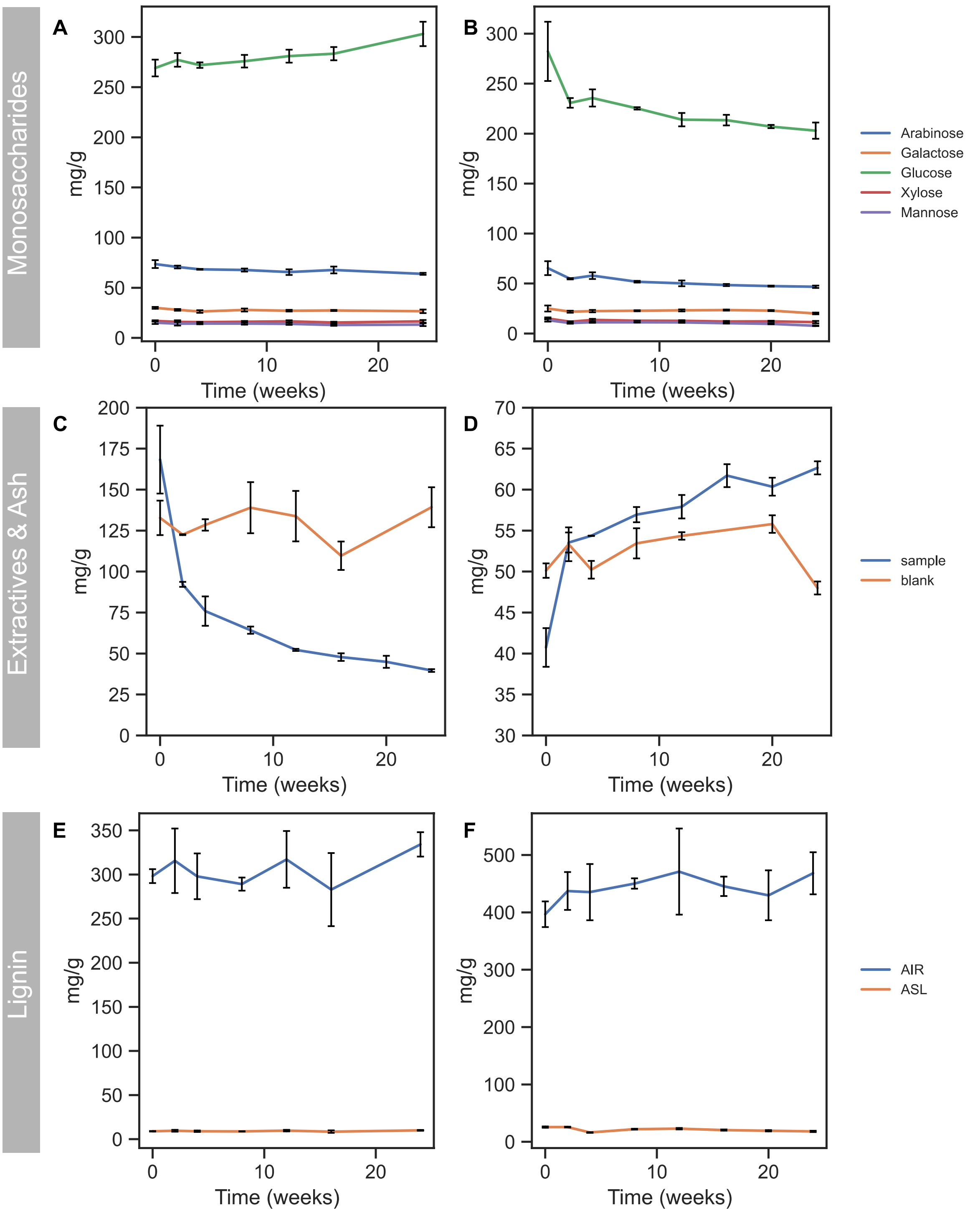

**Figure S1. Chemical analyses of lignin, carbohydrates, and extractives.** Effect of microbial growth on spruce bark on uninoculated control and monosaccharide composition after sulfuric acid hydrolysis for A) uninoculated control, and B) biotic sample. Total acetone extract C) and D) ash content. and effect on acid insoluble residue (AIR) and acid soluble lignin (ASL) for E) uninoculated control, and F) biotic sample. Mean and standard deviations are based upon duplicate biological experiments and two technical replicates except extractive and ash measurements which are based on biological triplicate experiments.

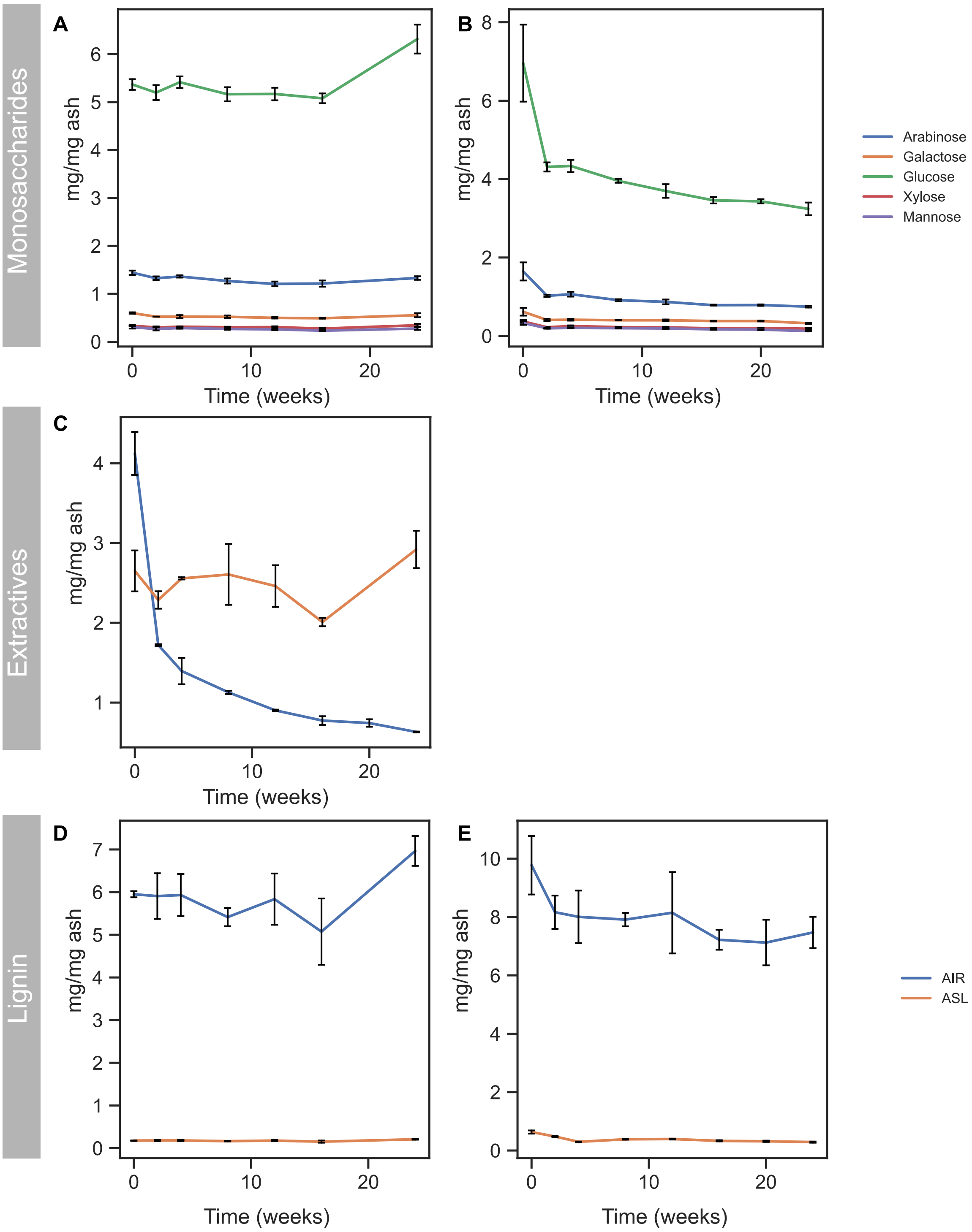

**Figure S2. Chemical analyses (as in Fig. S2) normalized against the ash content.** Acid insoluble residue (AIR) and acid soluble lignin (ASL) in the A) uninoculated control and B) biotic sample. Monosaccharide composition after sulfuric acid hydrolysis in the C) uninoculated control and D) biotic sample. Total acetone extract in E). Mean and standard deviation are based upon duplicate biological experiments and two technical replicates for all experiments except extractive measurements which are based on biological triplicate experiments.

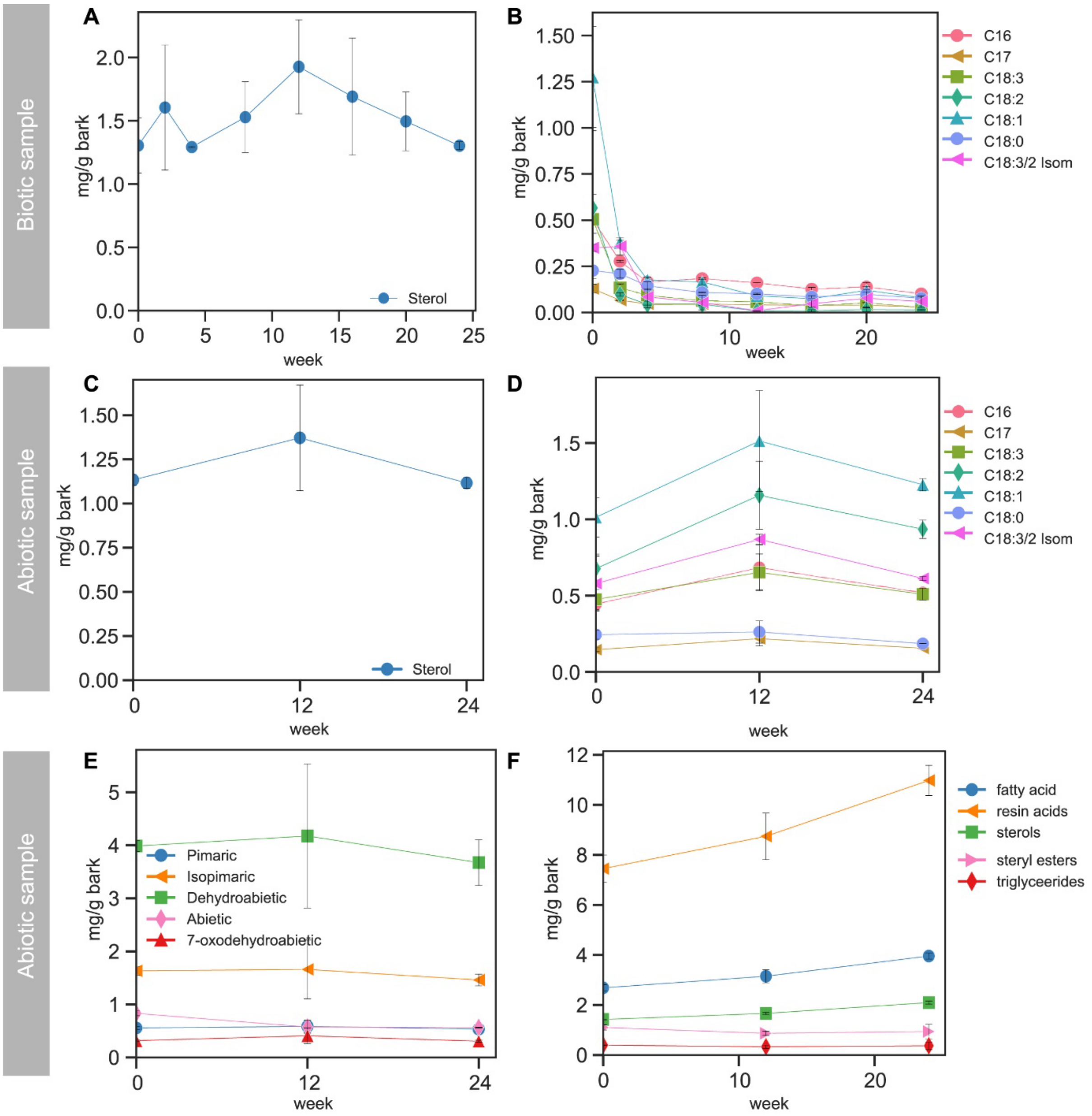

**Figure S3. Changes in the extractive groups and compounds in the biotic and the abiotic (blank) sample.** A) Unidentified sterol at RT=24.33 in the biotic sample. B) Fatty acids in the biotic sample. C) Extractive groups in the abiotic sample (blank). D) Unidentified sterol at RT=24.33 in the abiotic sample. D) Resin acids in the abiotic sample. E) Fatty acids in the abiotic sample. Mean and standard deviations are based upon duplicate biological experiments.

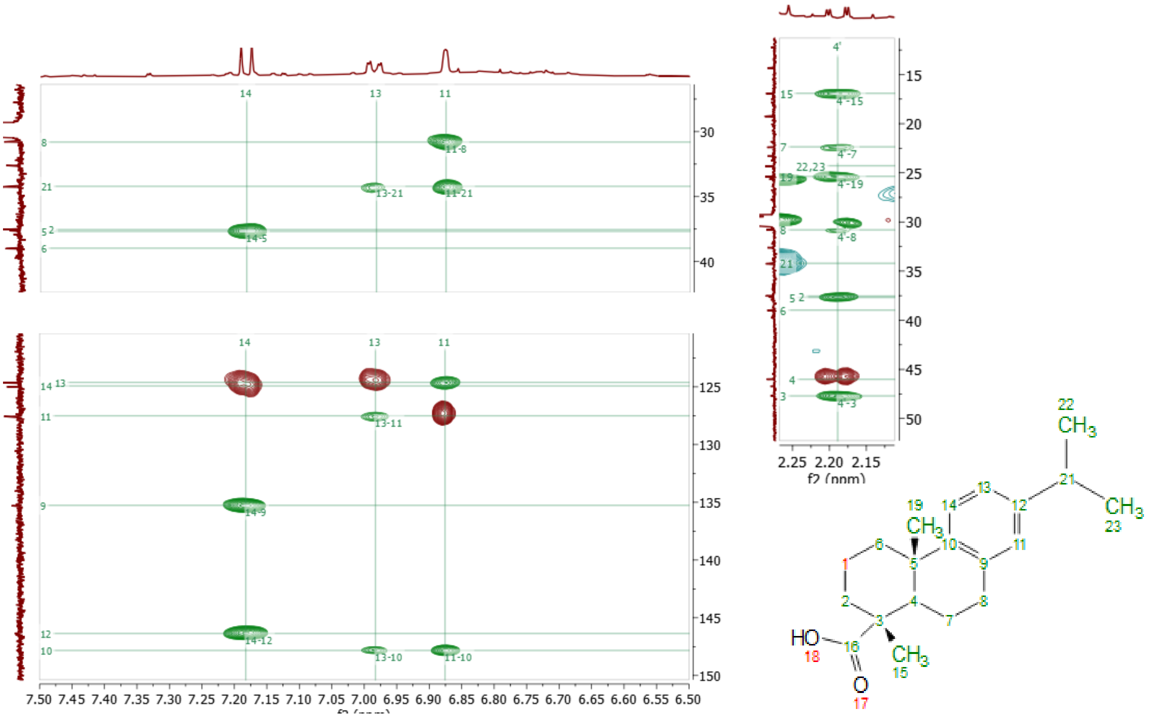

**Figure S4. Structure elucidation of dehydroabietic acid in spruce bark extract from the two-week degraded sample using 2D-NMR.** Important cross-signals in HSQC (red) and HMBC (green) aromatic region with assignment of dehydroabietic acid. The proton 4 showed several correlations confirming this structure, including the correlations to carbons 2, 3, 7, 8, 15, 16, and 19. Methyl protons on 22/23 show clear correlations to the aromatic carbon 12 whereas the methyl protons 19 show correlations to the aromatic carbon 10 and the methyl protons 15 have correlation to the carboxylic acid 16, confirming the presence of this group.

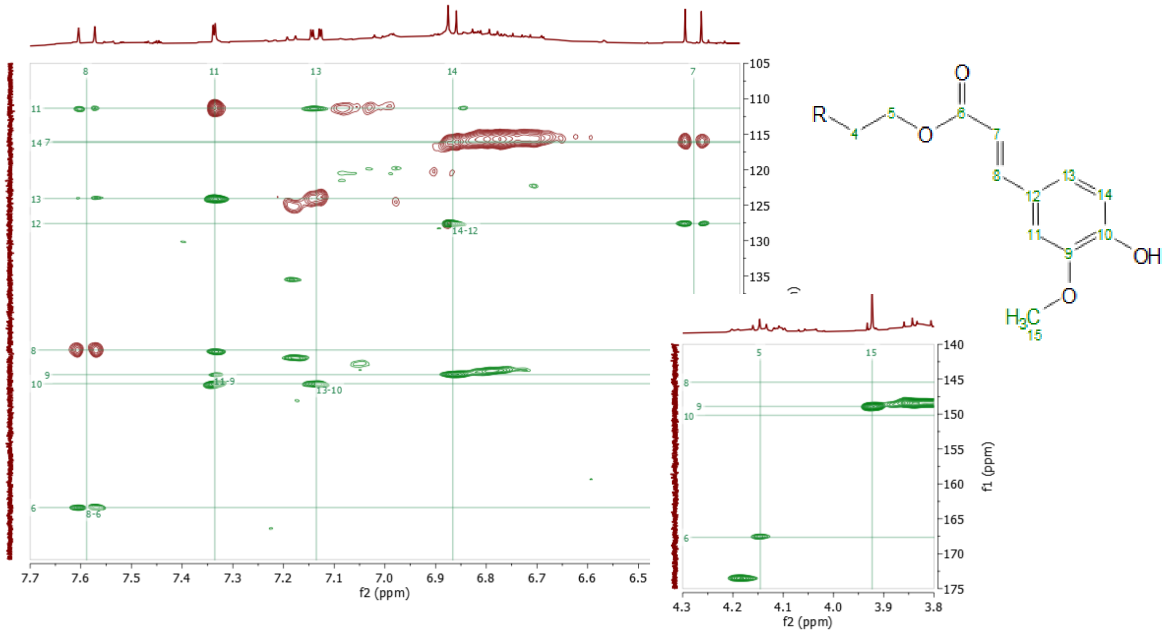

**Figure S5. Structure elucidation of ferulic ester compound in spruce bark extract from the 24-week degraded sample using 2D-NMR.** Important cross-signals in HSQC (red) and HMBC (green) aromatic region with assignment of a ferulic ester compound. Inset show a correlation between aromatic ring carbon 9 to methoxy group proton 15, as well as a correlation between the ester carbon 6 towards a proton 5. Clear cross signals originating from the aromatic ring as well as cross signals originating from a double bond can be seen in the spectrum. For the double bond, the J-couplings of 16 Hz corresponded well to those obtained from a double bond in trans figuration[8]. From the HMBC cross signals connecting the aromatic ring and the double bond could be seen. Most notably, the proton on 8 have cross-correlations in the HMBC to the aromatic carbons 11 and 13. In addition to this, the 8 proton also have a correlation to an ester carbon 6. In the inset of the figure, a correlation between the ester carbon 6 towards a proton 5 can be seen indicating that this is a ferulic ester compound rather than a ferulic acid.

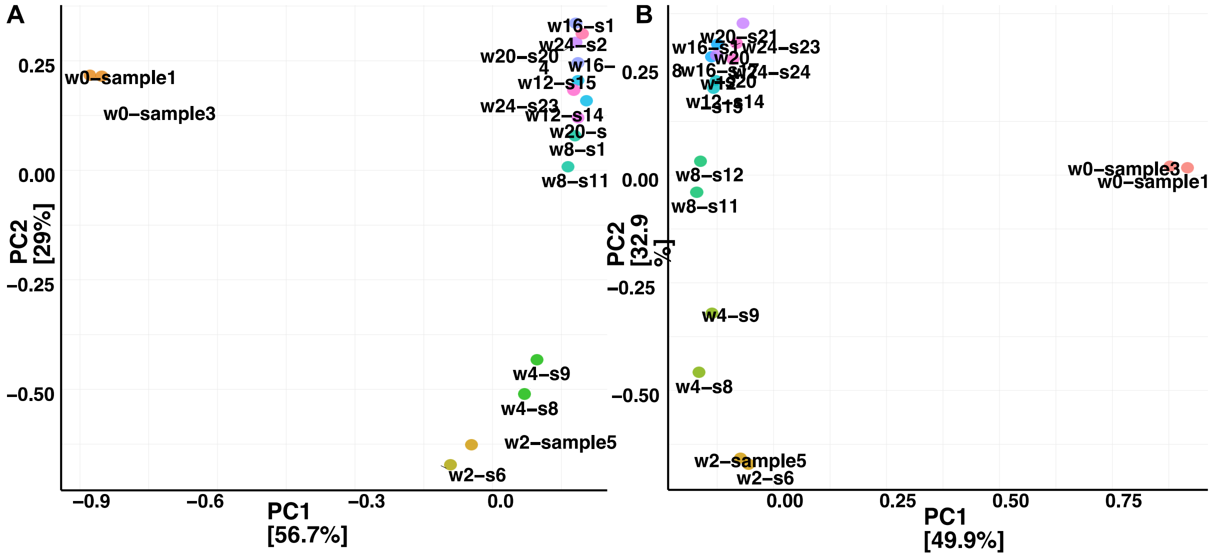

**Figure S6. Principle component analysis (PCA).** Identification of samples with similar microbial community was done using multivariate statistics, to illustrate sample similarities. Each point represents a community in a specific sample and is colored by the week sampled, and good agreement between biological replicates was confirmed by principal component analysis (PCA) A) bacteria B) fungi.

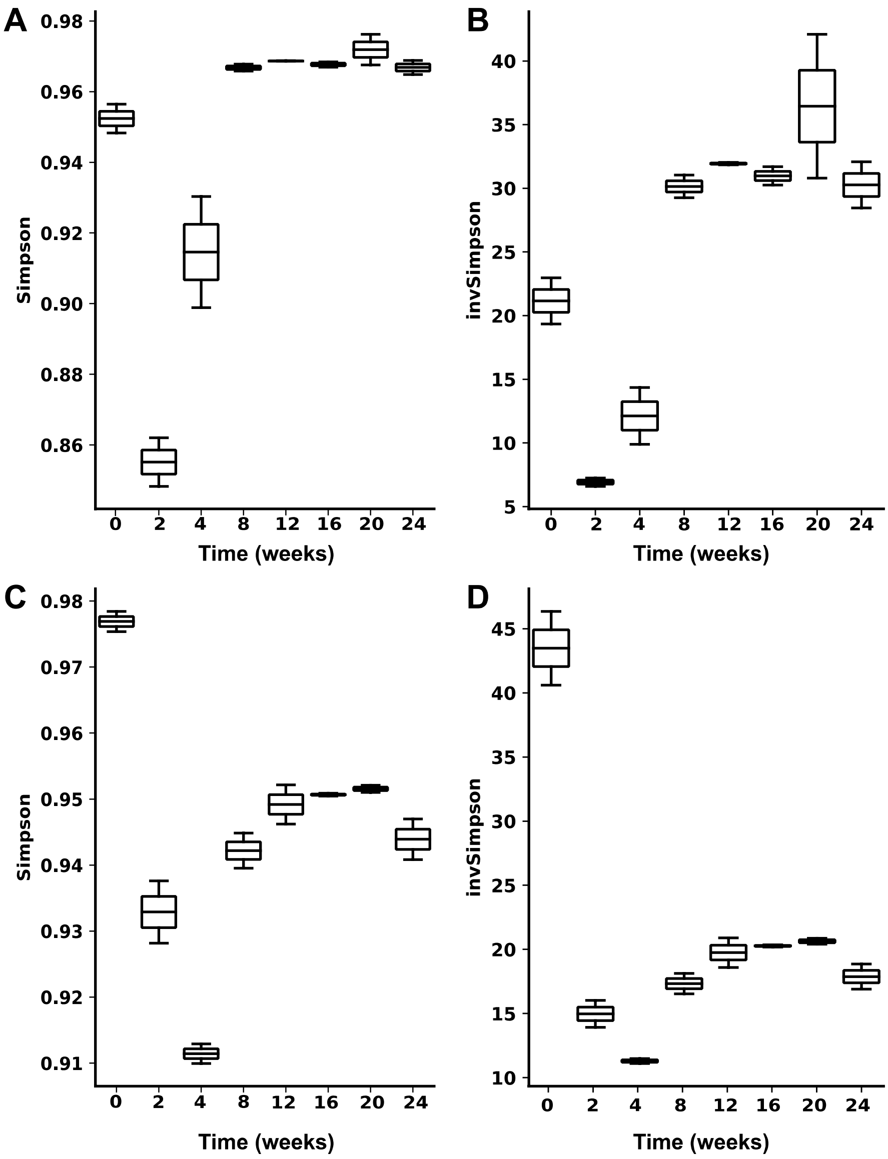

**Figure S7. Alpha-diversity index of microorganisms growing on spruce bark over time.** For bacteria A) Simpson index, and B) invSimpson index. For fungi C) Simpson index, D) invSimpson index. Mean and standard deviations are based upon duplicate biological experiments.

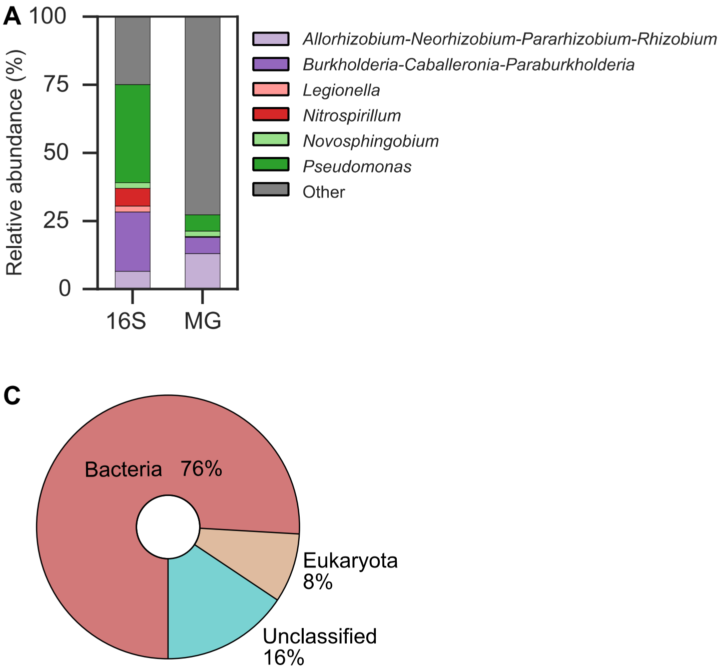

**Figure S8. Microbial taxonomic composition after two weeks of spruce bark degradation.** A) Relative phyla, B) class abundances based on 16S rRNA gene target sequencing (16S) and whole metagenome (MG) reads. C) comparison of fungal, bacterial and unclassified abundance in sample.

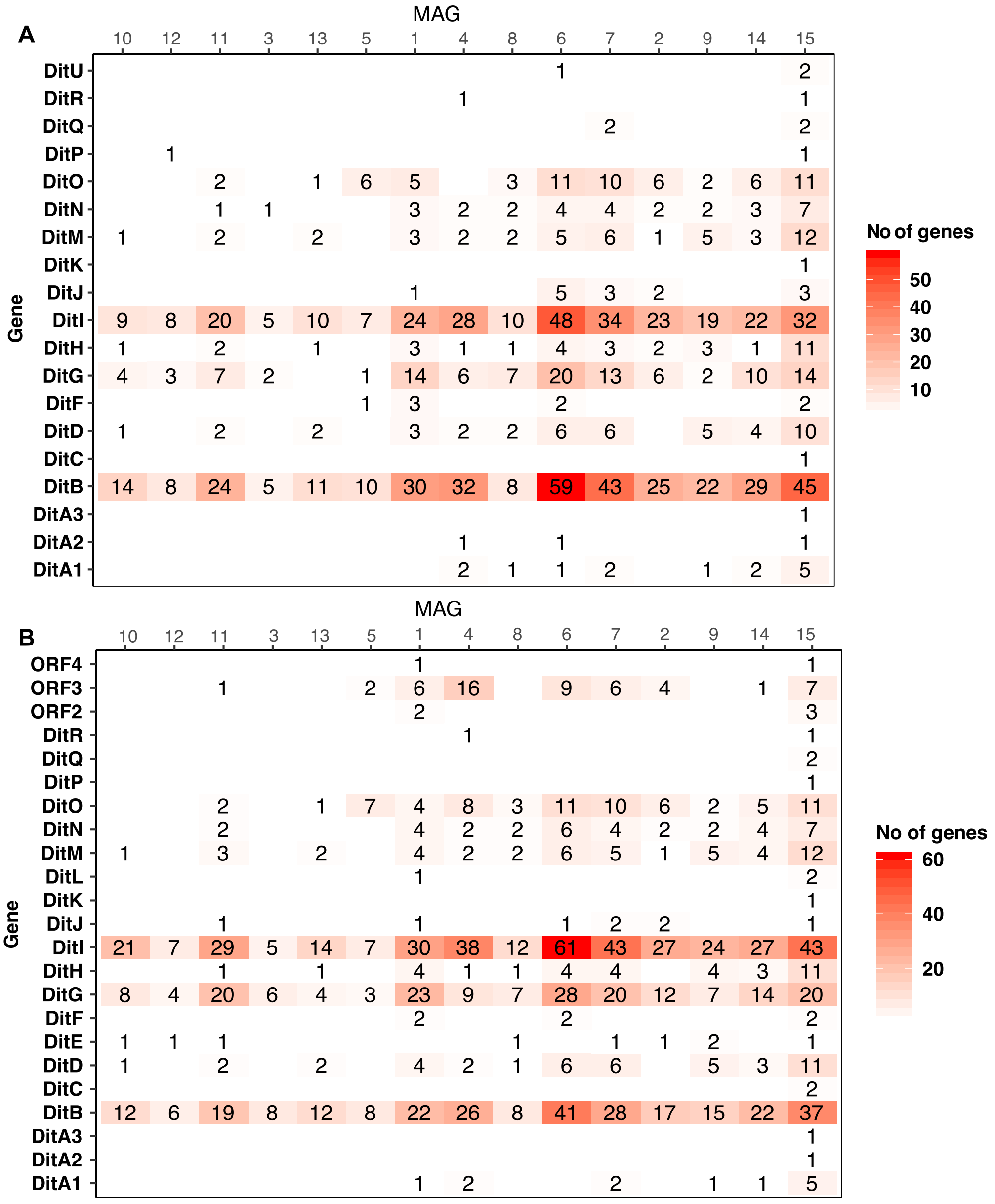

**Figure S9. Abundance of *dit* cluster proteins in MAGs.** BLAST analysis of MAGs derived from two-week sample of degraded spruce bark. Using either proteins A) *Paraburkholderia xenovorans* LB400 B) *Pseudomonas abietaniphila* BKME-9 as a query sequence.

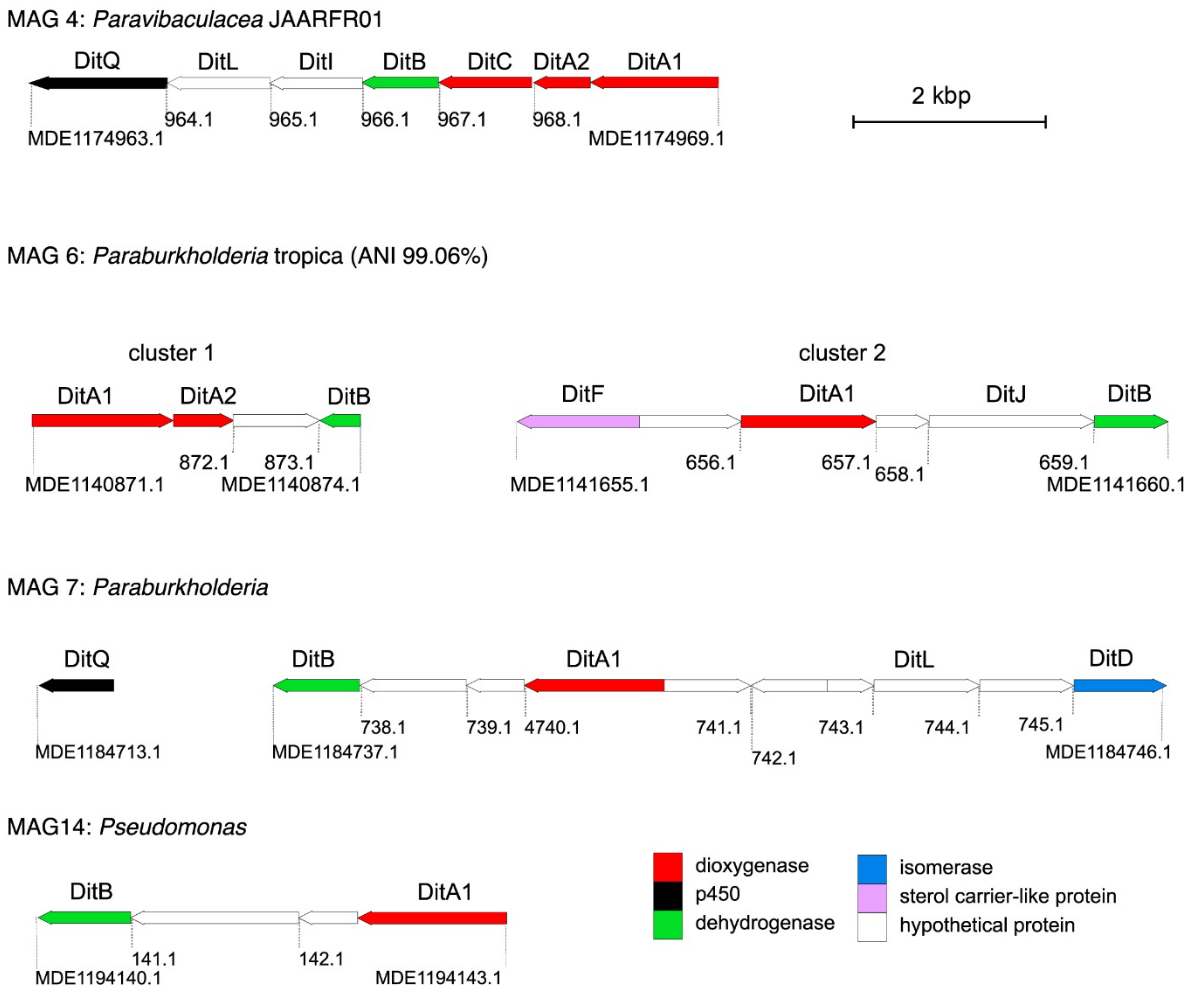

**Figure S10. Genetic organization of putative *dit* gene clusters, or *dit*-cluster-like fragments, in the MAGs.** Genes encoding proteins of predicted functions (BLAST) are color coded, and locus tags are indicated below the start and end of each gene cluster.

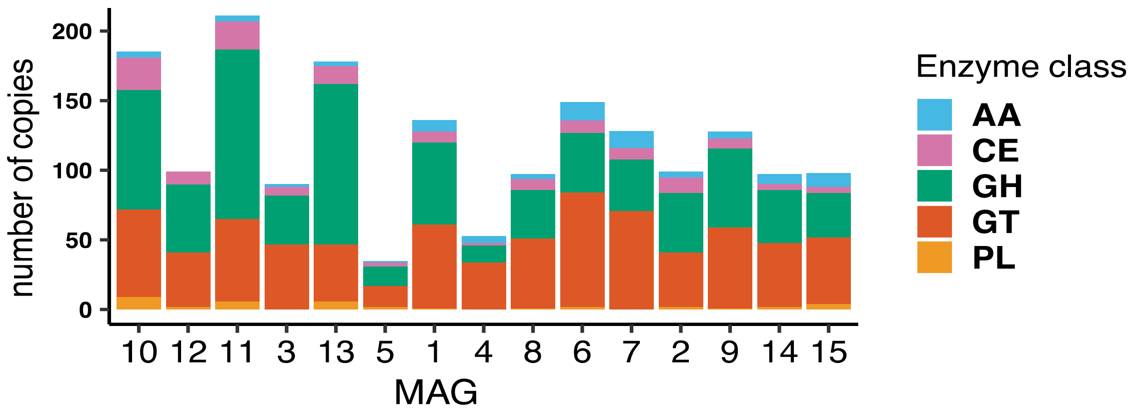

**Figure S11. CAZyme analysis of the MAGs.** The different CAZy classes are colour coded, with auxiliary activity (AA) in blue, carbohydrate esterase (CE) in purple, glycoside hydrolase (GH) in green, glycosyltransferase (GT) in red, and polysaccharide lyase (PL) in orange.

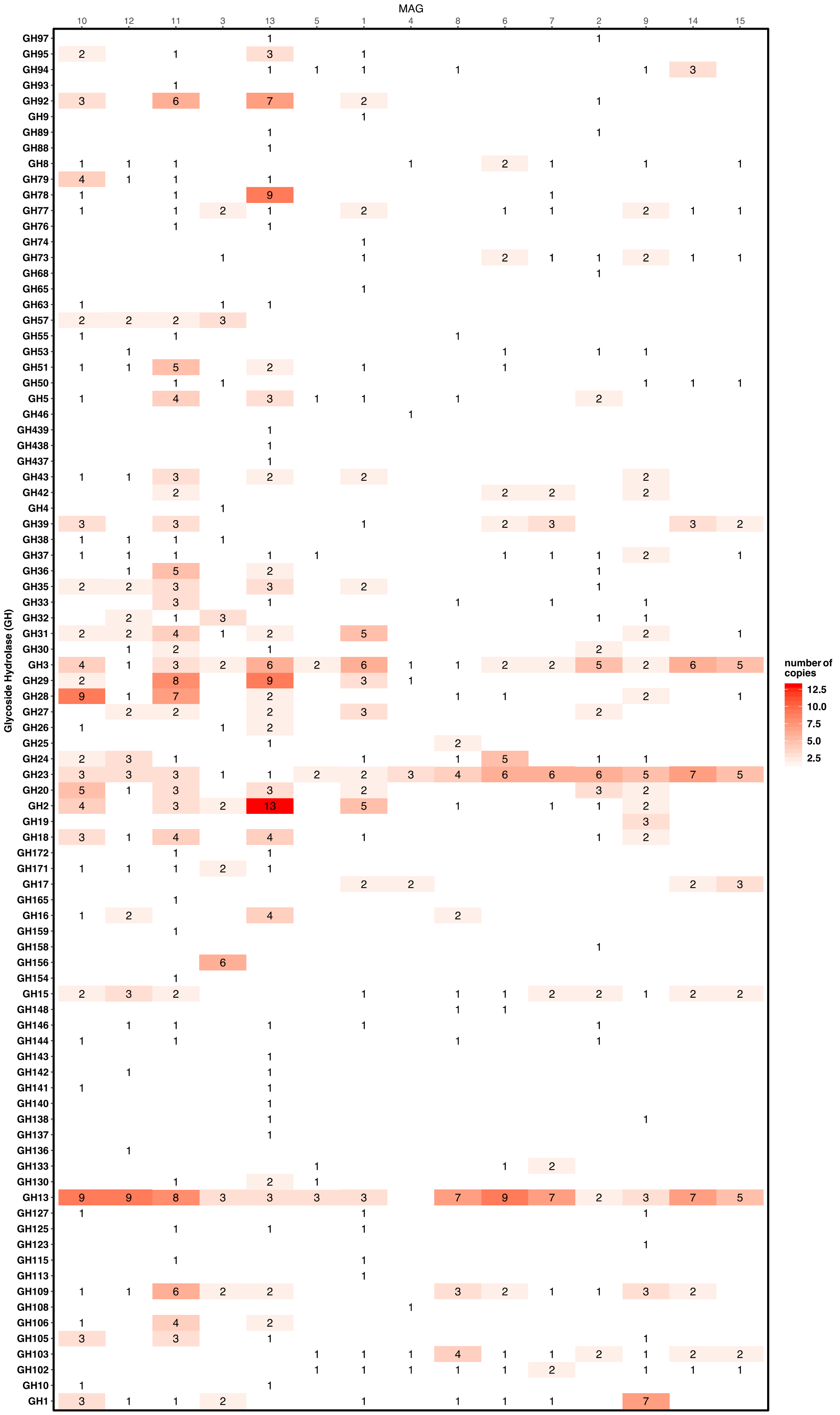

MAG
10 12 11 3 13 5 1 4 8 6 7 2 9 14 15
Glycoside Hydrolase (GH)
GH97 GH95 GH94 GH93 GH92 GH9 GH89 GH88 GH8 GH79 GH78 GH77 GH76 GH74 GH73 GH68 GH65 GH63 GH57 GH55 GH53 GH51 GH50 GH5 GH46 GH439 GH438 GH437 GH43 GH42 GH4 GH39 GH38 GH37 GH36 GH35 GH33 GH32 GH31 GH30 GH3 GH29 GH28 GH27 GH26 GH25 GH24 GH23 GH20 GH2 GH19 GH18 GH172 GH171 GH17 GH165 GH16 GH159 GH158 GH156 GH154 GH15 GH148 GH146 GH144 GH143 GH142 GH141 GH140 GH138 GH137 GH136 GH133 GH130 GH13 GH127 GH125 GH123 GH115 GH113 GH109 GH108 GH106 GH105 GH103 GH102 GH10 GH1
number of copies
12.5
10.0
7.5
5.0
2.5

**Figure S12.** Number of predicted copies and CAZy family membership of MAG glycoside hydrolases (GH).

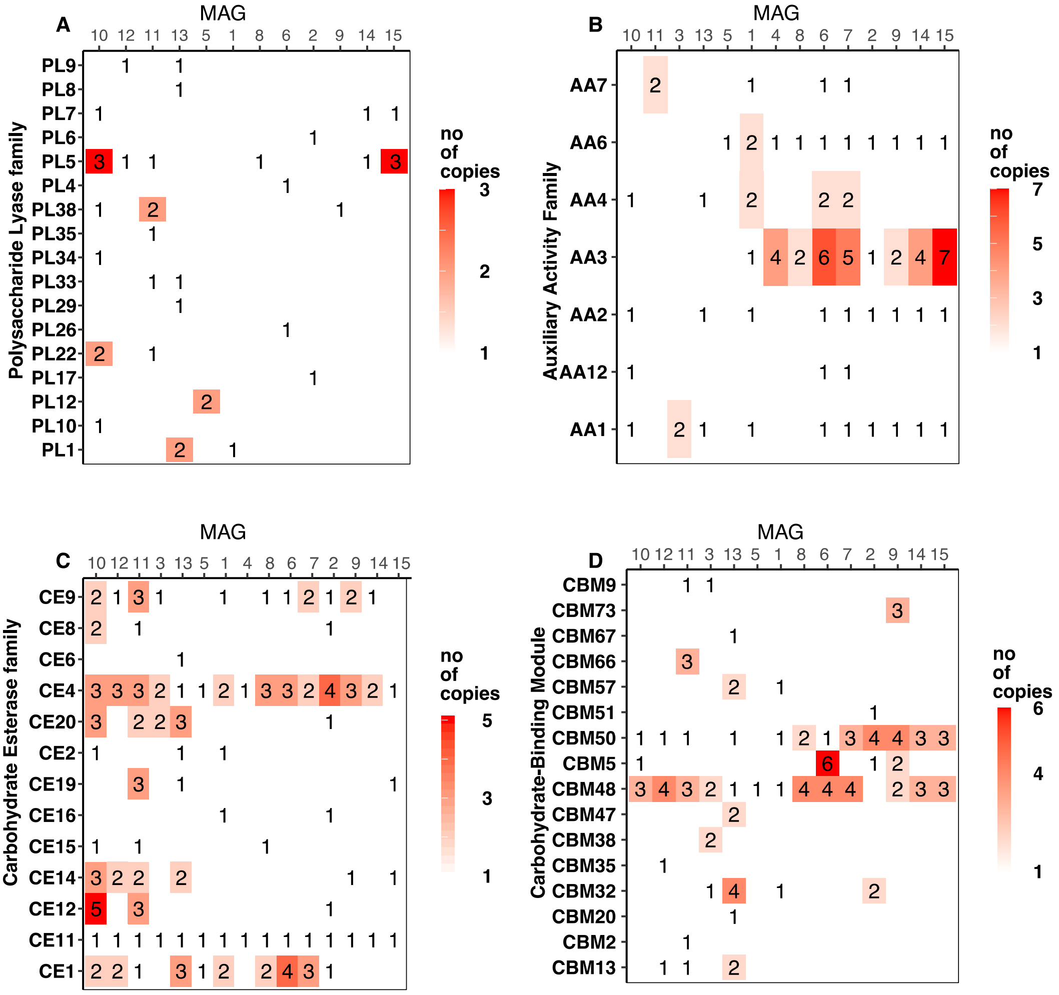

**Figure S13.** Number of predicted copies and CAZy family membership in the MAGs for A) polysaccharide lyases, B) auxiliary activities, C) carbohydrate esterases, and D) carbohydrate binding modules (CBMs).

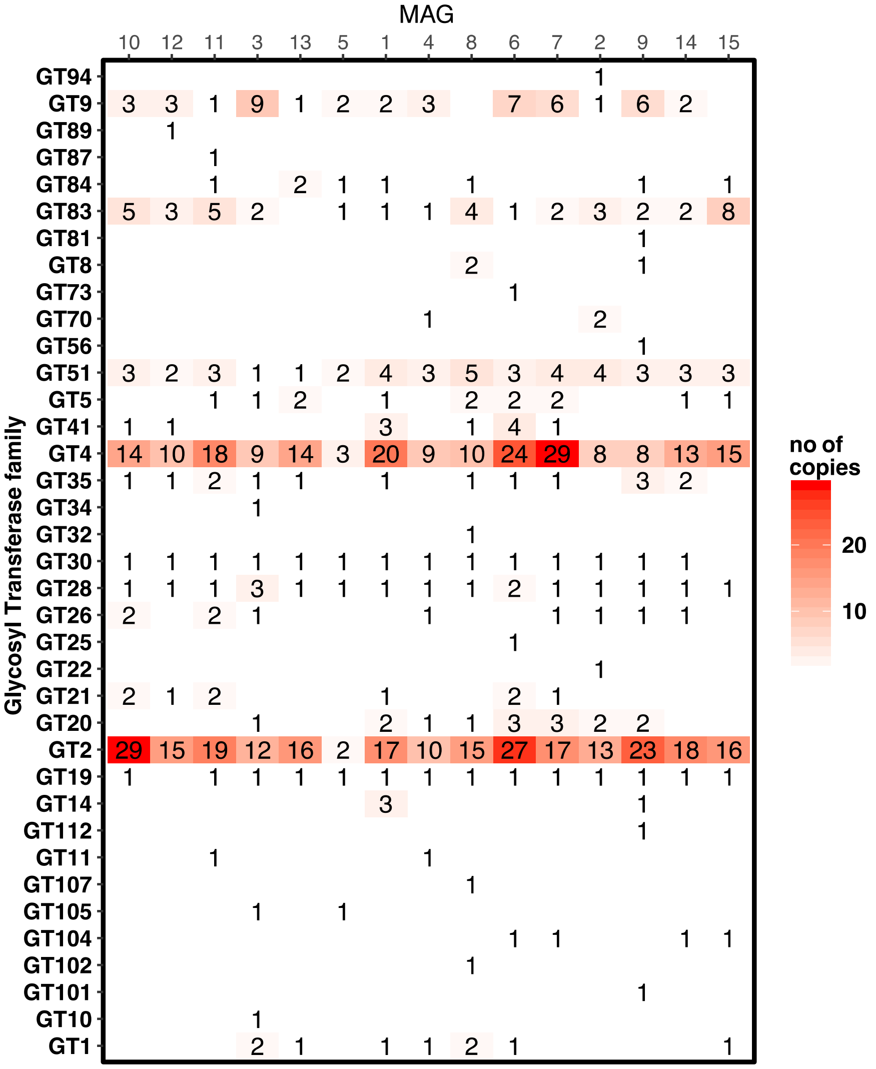

**Figure S14.** Number of predicted copies and CAZy family membership of MAG glycosyltransferases (GTs).

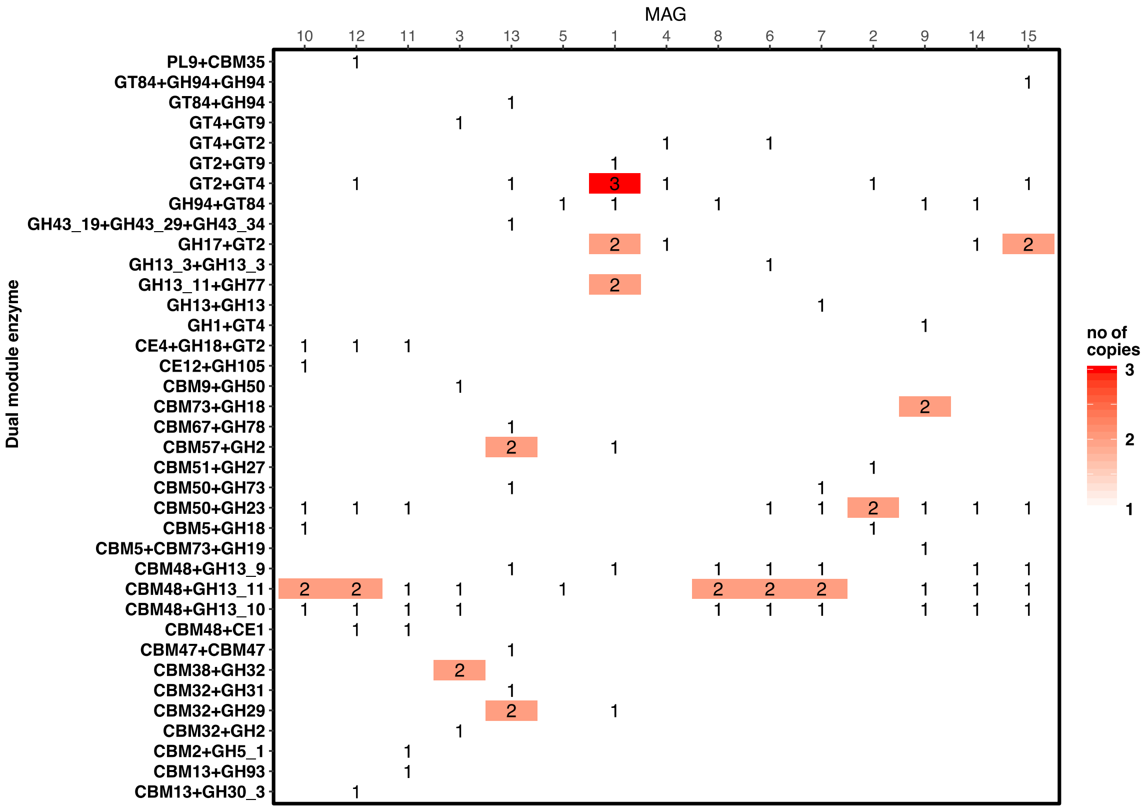

**Figure S15.** Multimodular CAZymes identified in the MAGs, and their domain order.

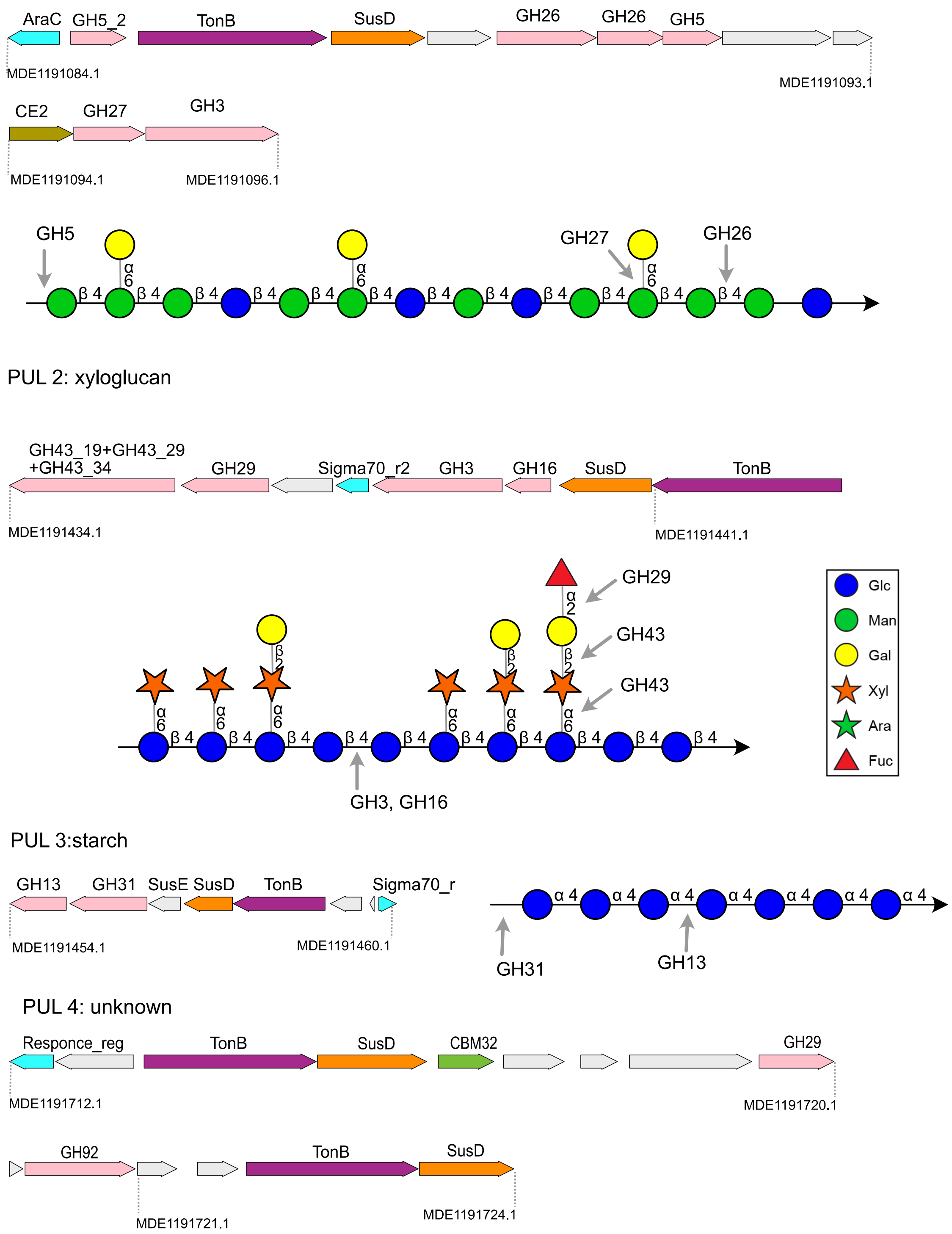

**Figure S16a. Putative Polysaccharide Utilization Loci (PULs) identified in MAG13.** The PUL-encoded genes are colored according to putative function, and gray arrows indicate both putatively annotated proteins and proteins of unknown function. Representative polysaccharide structures are shown close to each identified PUL.

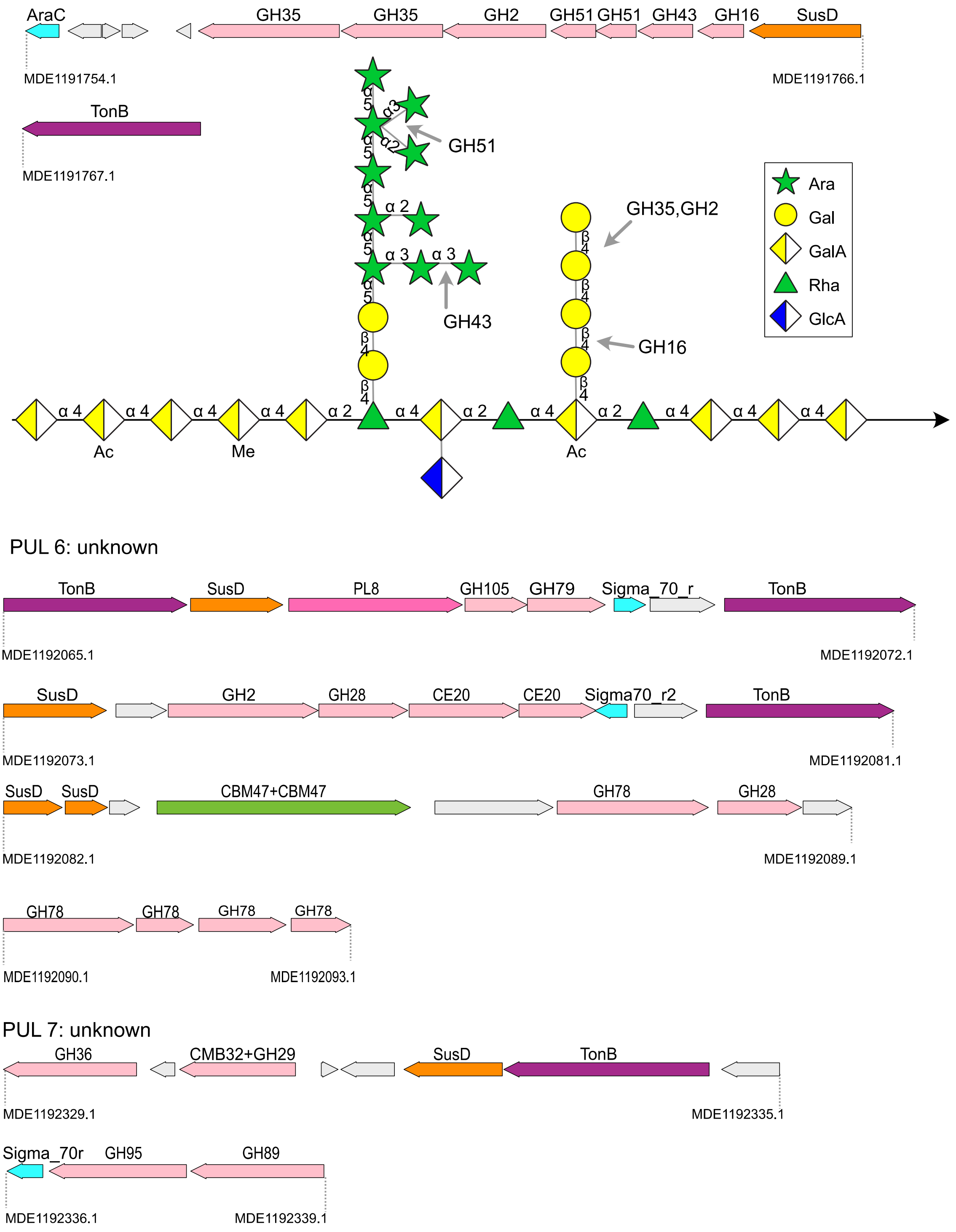

**Figure S16b. Putative Polysaccharide Utilization Loci (PULs) identified in MAG13.** The PUL-encoded genes are colored according to putative function, and gray arrows indicate both putatively annotated proteins and proteins of unknown function. Representative polysaccharide structures are shown close to each identified PUL.

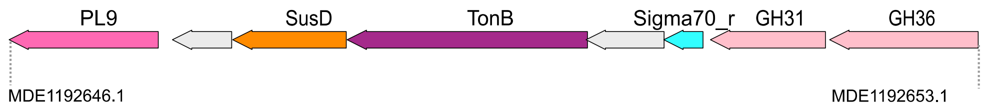

PUL 9:unknown

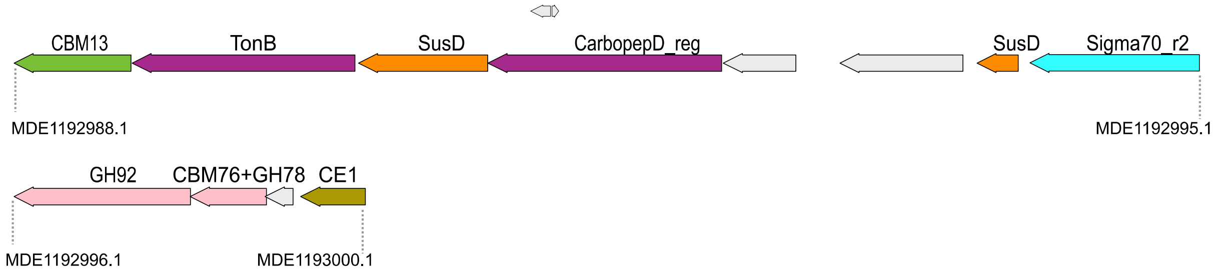

PUL 10: RG-II

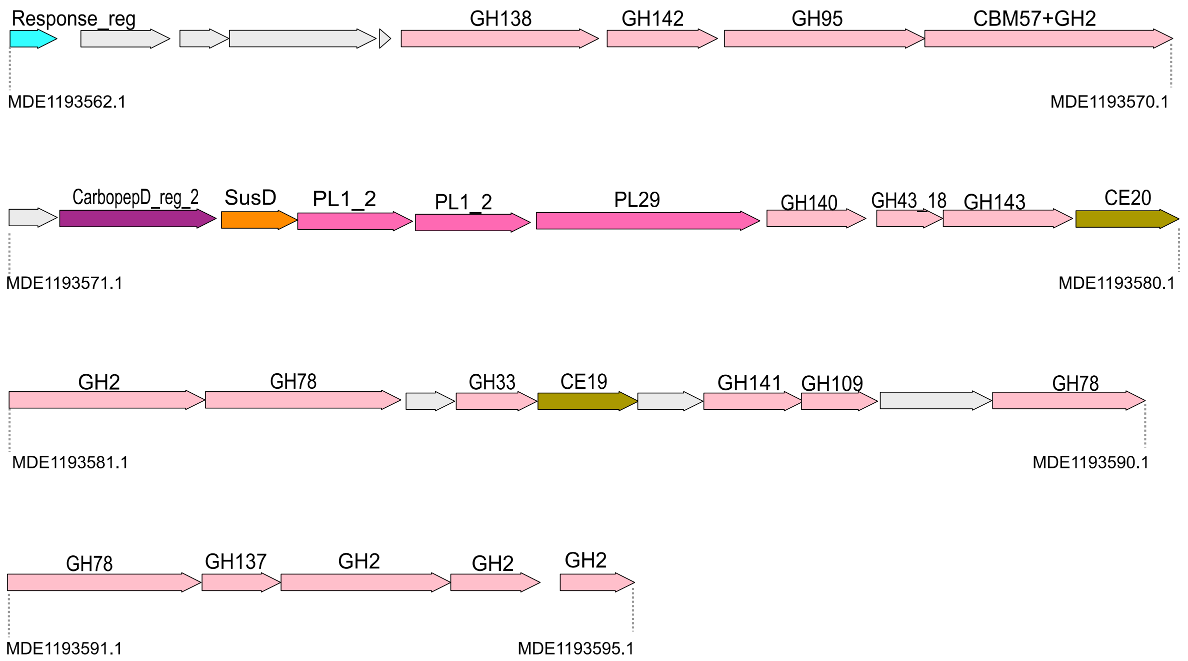

**Figure S16c. Putative Polysaccharide Utilization Loci (PULs) identified in MAG13.** The PUL-encoded genes are colored according to putative function, and gray arrows indicate both putatively annotated proteins and proteins of unknown function. Representative polysaccharide structures are shown close to each identified PUL.

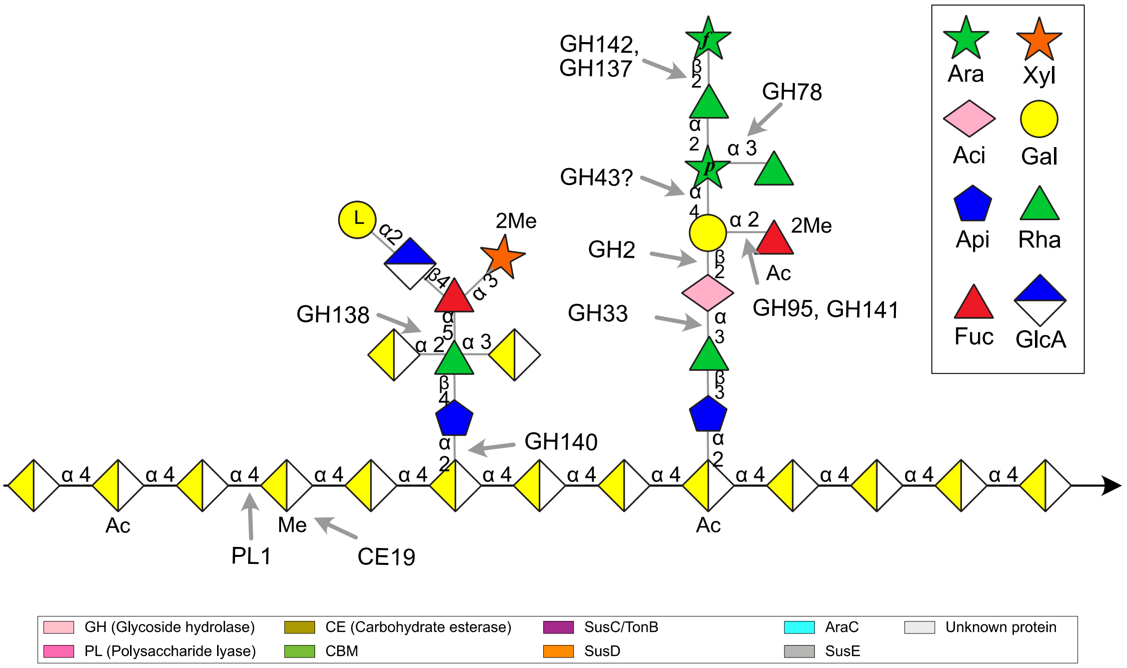

**Figure S16d. Putative Polysaccharide Utilization Loci (PULs) identified in MAG13.** The PUL-encoded genes are colored according to putative function, and gray arrows indicate both putatively annotated proteins and proteins of unknown function. Representative polysaccharide structures are shown close to each identified PUL.

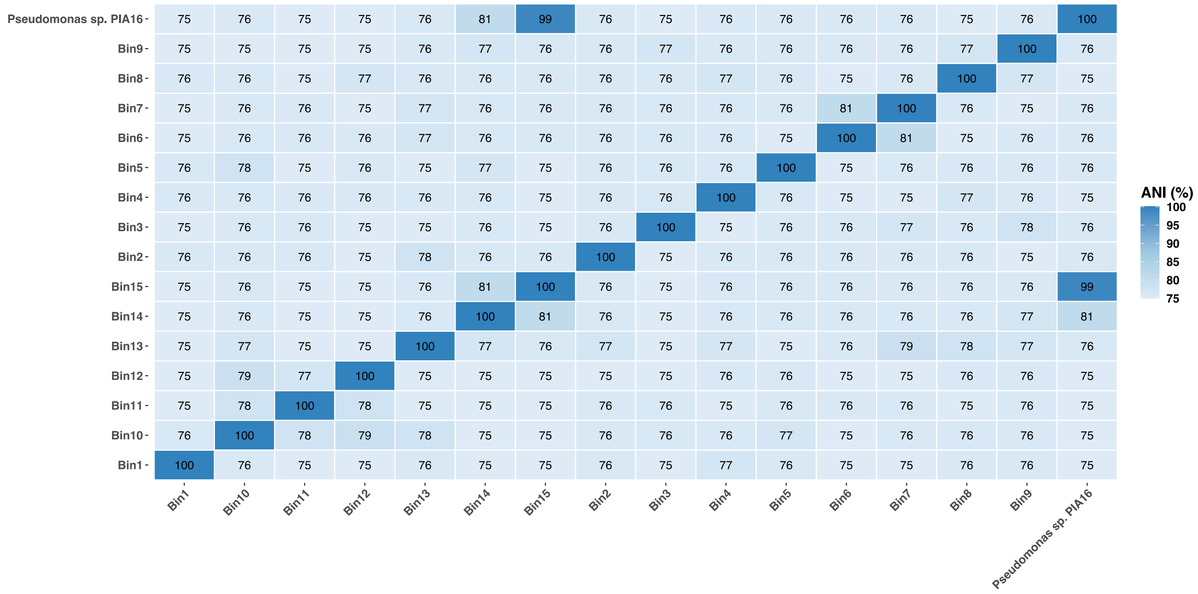

**Figure S17.** Average nucleotide identity (ANI) between the extracted MAGs (Bins) from the two-week bark metagenome assembly and the isolate *Pseudomonas sp*. PIA16 (*Pseudomonas abieticivorans* sp. nov).

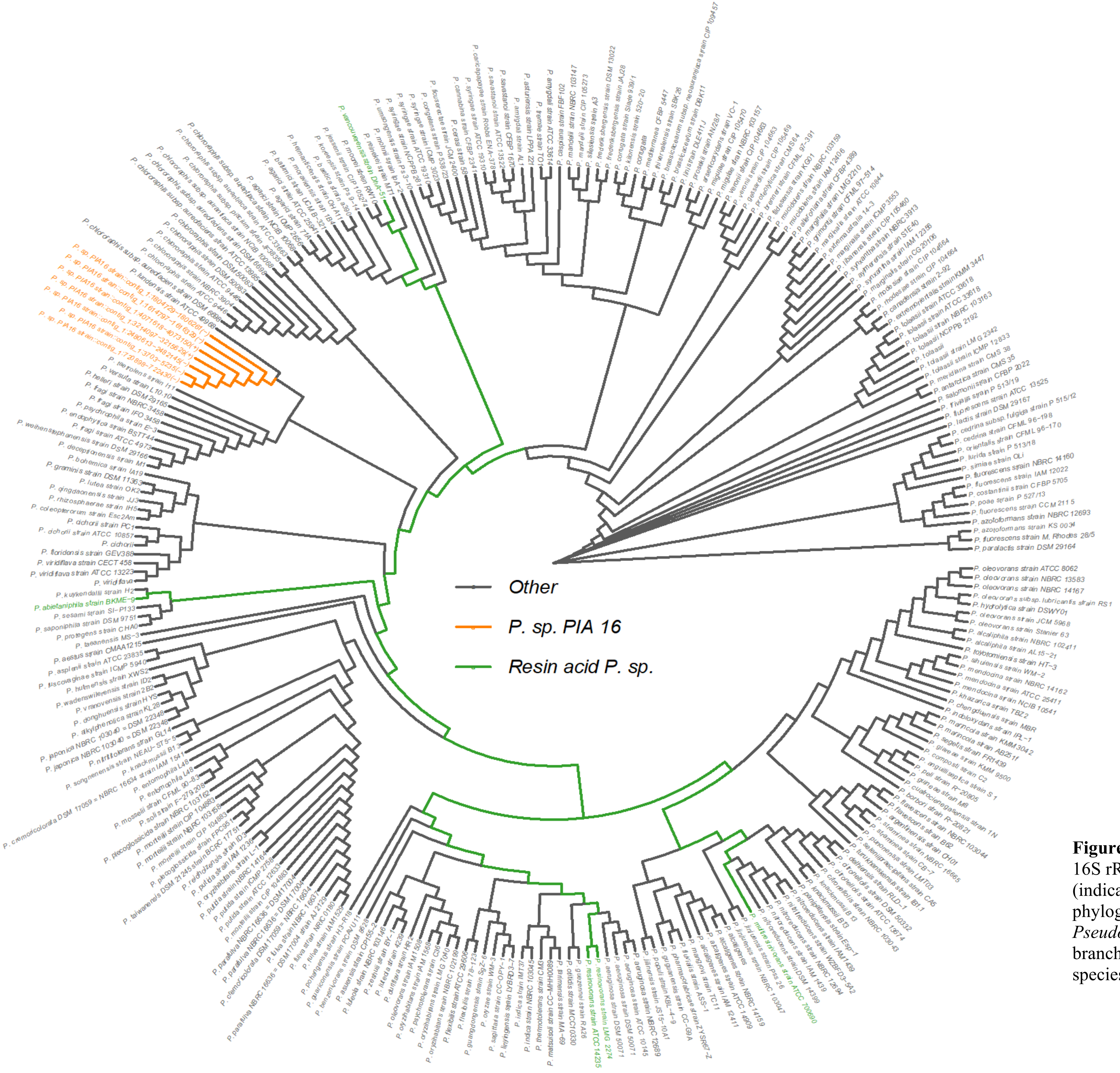

**Figure S18.** Phylogenetic tree based on the 16S rRNA gene of *P. abieticivorans* (indicated in orange as *P. sp.* PIA 16) and phylogenetically close members of *Pseudomonas* from NCBI: GenBank. Green branches represent known *Pseudomonas* species able to metabolize resin acids.

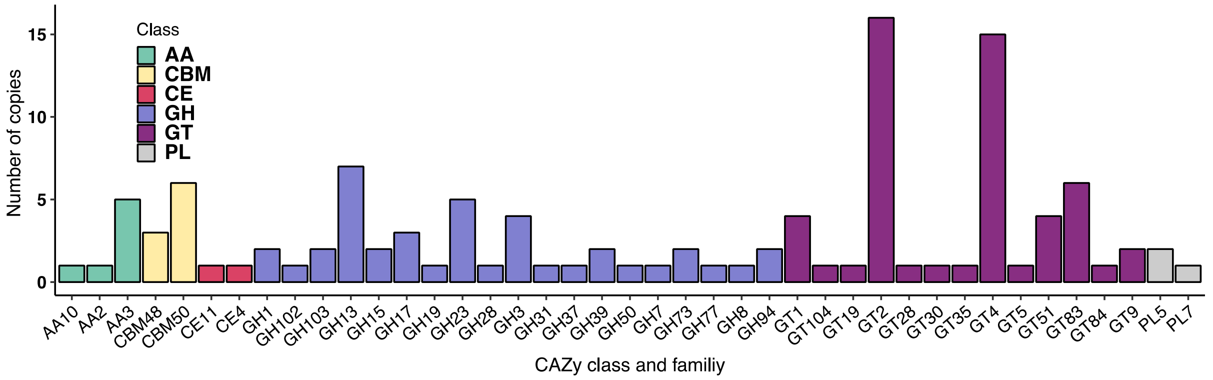

**Figure S19.** CAZy families and copy numbers putatively encoded by *Pseudomonas abieticivorans*. AA – auxiliary activity, CBM – carbohydrate binding module, CE – carbohydrate esterase, GH – glycoside hydrolase, GT – glycosyltransferase, PL – polysaccharide lyase.

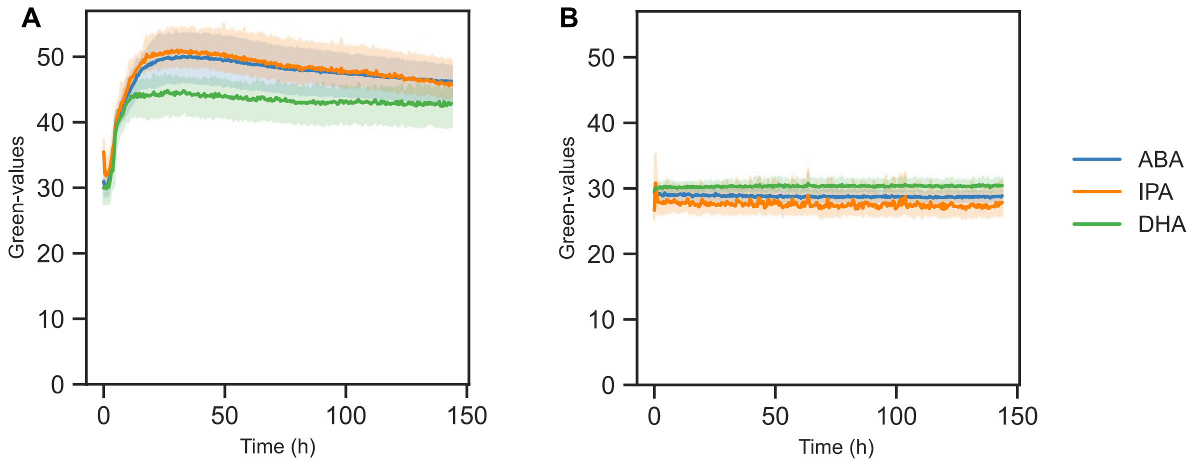

**Figure S20.** Growth profiles of *P. abieticivorans* on abietic acid (AA), isopimaric acid (IPA), and dehydroabietic acid. Growth Profiler Green Values corresponding to growth based on pixel counts. A) of *P. abieticivorans* B) non-inoculated blank sample. The growth profiles are shown as averages of biological triplicate experiments in bold colored lines and lighter colors are values from all experiments.

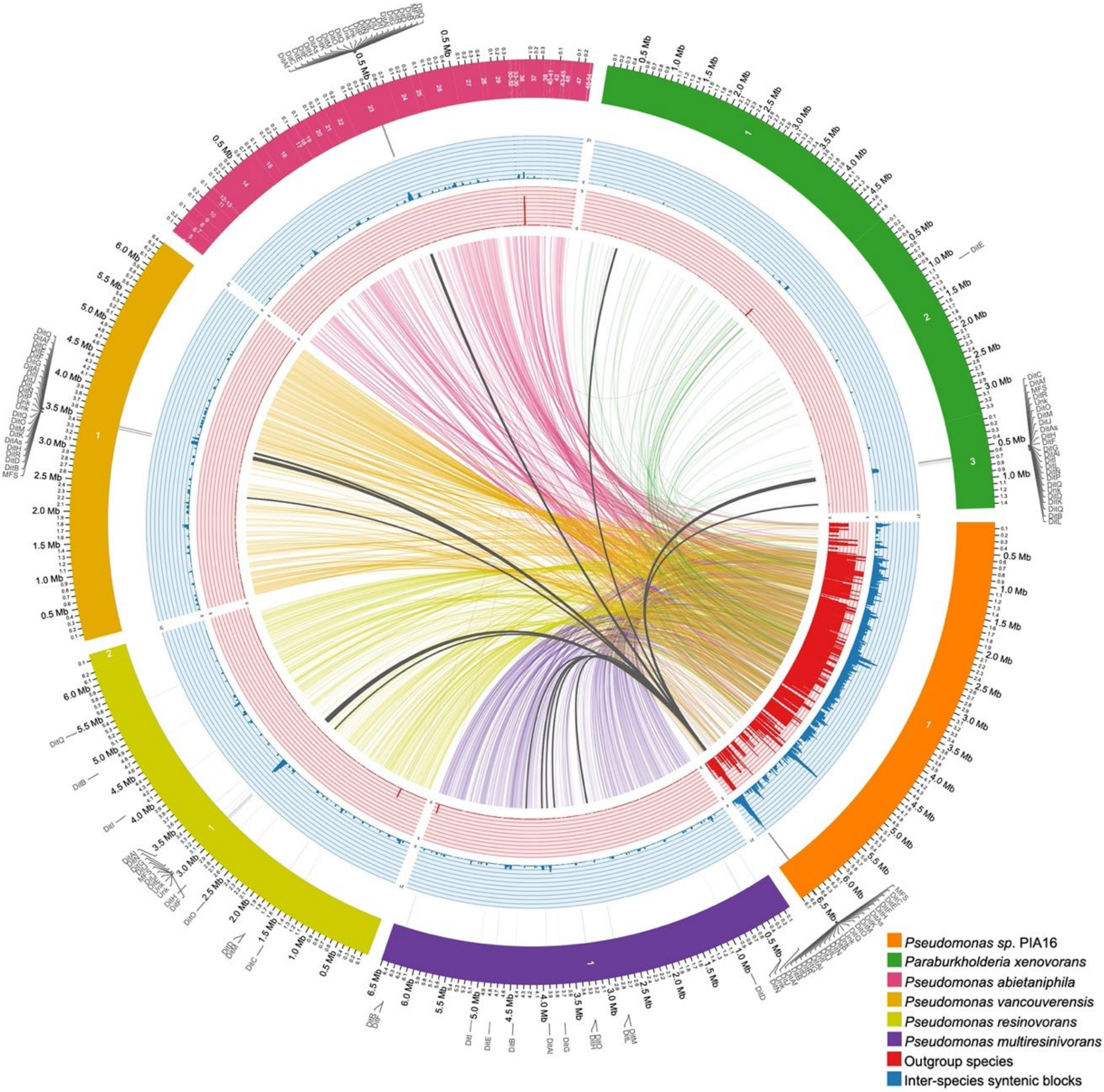

**Figure S21.** Collinear analysis of five known resin acid utilizing strains and *P. abieticivorans* (sp. PIA16). Numbers inside the ideograms represent chromosome number, plasmid (two in *P. resinovorans*), or contigs (in *P. abietaniphila*). Links between genomes represents coding sequences in synteny blocks. Dark gray links indicate *dit* cluster genes.
